# Improving biodiversity in Central and Eastern European domestic gardens needs regionally scaled strategies

**DOI:** 10.1101/2024.07.12.603327

**Authors:** Zsófia Varga-Szilay, Arvīds Barševskis, Klára Benedek, Danilo Bevk, Agata Jojczyk, Anton Krištín, Jana Růžičková, Lucija Šerić Jelaska, Eve Veromann, Silva Vilumets, Kinga Gabriela Fetykó, Gergely Szövényi, Gábor Pozsgai

**Affiliations:** Doctoral School of Biology, Institute of Biology, ELTE Eötvös Loránd University, Pázmány Péter sétány 1/C, 1117, Budapest, Hungary; Coleopterological Research Center, Institute of Life Sciences and Technology, Daugavpils University, Vienības str. 13, 5401, Daugavpils, Latvia; Department of Horticulture, Faculty of Technical and Human Sciences, Sapientia Hungarian University of Transylvania, Calea Sighișoarei nr. 2, 540485, Târgu-Mureș, Romania; Department of Organisms and Ecosystems Research, National Institute of Biology, Vecna pot 121, 1000, Ljubljana, Slovenia; Department of Landscape Art, Institute of Environmental Engineering, Warsaw University of Life Science, Nowoursynowska str. 166., 02-787, Warsaw, Poland; Institute of Forest Ecology Slovak Academy of Sciences, Ľ. Štúra 2, 960 53, Zvolen, Slovakia; HUN-REN-ELTE-MTM Integrative Ecology Research Group, ELTE Eötvös Loránd University, Pázmány Péter sétány 1/C, 1117, Budapest, Hungary; Department of Systematic Zoology and Ecology, ELTE Eötvös Loránd University, Pázmány Péter sétány 1/C, 1117, Budapest, Hungary; Department of Biology, Faculty of Science University of Zagreb, Rooseveltov trg 6, 10000, Zagreb, Croatia; Chair of Plant Health, Institute of Agricultural and Environmental Sciences, Estonian University of Life Sciences, Fr. R. Kreutzwaldi 1, 51006, Tartu, Estonia; Independent researcher, 6000, Kecskemét, Hungary; Ce3C – Centre for Ecology, Evolution and Environmental Changes, Azorean Biodiversity Group, CHANGE – Global Change and Sustainability Institute, University of the Azores, Faculty of AgricuGltural Sciences and Environment, Rua Capitão João D’Ávila, 9700-042, Angra do Heroísmo, Terceira, Açores, Portugal

**Keywords:** rural-urban gradient, urban ecosystems, environmental consciousness, sustainable gardening, environmental sensitivity, urbanisation

## Abstract

Amid ongoing urbanisation and increasing anthropogenic activities, domestic gardens, while they cannot replace undisturbed habitats, play a valuable role in enhancing urban biodiversity by supplementing green areas and improving landscape connectivity. Moreover, biodiversity-friendly gardens also improve human well-being and foster connections between nature and people. To study these benefits, we distributed an online questionnaire between 2022 and 2023, and used a scoring system to evaluate the ecological value of gardens, garden owners’ motivations, and pesticide use habits. We used machine learning to explore how these indices interact and what sociodemographic factors drive them across nine Central and Eastern European (CEE) countries. Additionally, we explored the differences and similarities between the ecological values of gardens and gardening practices in building high biodiversity. Our findings reveal significant variability both between and within countries, across all three indices with Romania faring low and Czechia reaching high scores in all three indices. Domestic pesticide use was ubiquitous across all CEE and largely unaffected by sociodemographic factors. However, increased time spent with gardening was associated with reduced pesticide use and a greater potential for fostering high biodiversity. Garden owners over 55 tended to follow and uphold longstanding conventional practices and thus lowered both pesticide and garden index scores. The local-scale differences we observed emphasise the need for regionally tailored guidelines for biodiversity-friendly gardening over standardized regulations across Europe. Optimal strategies for effective environmental educational and community programs can be developed based on local biodiversity and the three indices used in this study. To maximize their impact and meet local needs, these programs should educate garden owners about the environmental and health effects of pesticides and offer comprehensive biodiversity-related information across all regions and social strata. This is particularly crucial in Central and Eastern Europe (CEE), where such initiatives are currently underrepresented.

**Highlights:**

- Domestic gardens can enhance biodiversity and support human well-being.
- Study conducted across nine Central and Eastern European countries.
- Significant variability both between and within countries in gardening practices.
- Improving biodiversity in domestic gardens needs regionally scaled strategies.
- Reducing pesticide use and increasing awareness are key.

## 1. Introduction

Built-up urban areas will cover 7% of the European Union by 2030 (Perpiña Castillo et al., 2019) and this ongoing urbanisation and increasing anthropogenic activities, in turn, contribute to biodiversity loss (Hagen et al., 2012) and the rapid decline of undisturbed ecosystems (Bengtsson et al., 2000). Most parts of Europe are already impoverished, and characterised by small natural areas, broken up by human intervention into a mosaic of agricultural lands and settlements (European Environment Agency, 2012; Eurostat, 2022b; Jongman, 2002; Perpiña Castillo et al., 2019). Thus, it is particularly important to halt, or at least reduce the ongoing degradation and losses of natural habitats in areas heavily modified by human activities, such as agricultural land and cities, and, as much as possible, maximise biodiversity. However, whilst conservation efforts for maintaining and preserving biodiversity have already been a part of many agricultural systems (Concepción et al., 2020; van Elsen, 2000), urban biodiversity rarely gains enough attention (Fischer et al., 2020; Ramalho & Hobbs, 2012). Although, the European Commission adopted a law to stop the loss of green urban spaces by 2030 and achieve a 10% a minimum tree canopy cover by 2050 (Nature Restoration Law, European Commission, 2024), the area of urban green spaces has only increased in some Southern and Western European cities (e.g. Pamplona/Iruña (Spain) and Hague (Netherlands)). From Eastern Europe, Czechia, Poland, and Romania are just about to catch up (European Environment Agency, 2023; Kabisch & Haase, 2013). Yet, in none of the European Union countries, could the amount of re-naturalised areas compensate for what has been lost for urbanisation (European Commission, 2022).

Urban green infrastructure covers an average of 42% of the city area according to the European Environment Agency (EEA) (European Environment Agency, 2022), implying that urban biodiversity can be substantially improved by developing green infrastructure (Baldock, 2020; Baldock et al., 2019; Heidt & Neef, 2008; Liang et al., 2023), such as diversifying city/urban parks and community gardens, rooftop gardens, establishing pollinator-friendly meadows, creating a network of corridors, and installing green roofs (Aguilera et al., 2019; Beninde et al., 2015; Lin et al., 2015; Seitz et al., 2022). Besides public places, however, private domestic gardens occupy an average of 16-36% of the total urban areas in Europe (Cameron et al., 2012; Goddard et al., 2010), and therefore the role of domestic gardens in preserving and supporting native biodiversity and their contribution to urban sustainability are becoming increasingly important (Fontaine et al., 2016; Hanson et al., 2021). Indeed, enhancing environmental quality in domestic gardens offers a great opportunity to improve human health and well-being (Camps-Calvet et al., 2016; Krols et al., 2022; Marques et al., 2021; Soga et al., 2017) and alleviate biodiversity loss simultaneously (Freeman et al., 2012; Li et al., 2023).

The biodiversity of domestic gardens varies along a wide spectrum (Home et al., 2019; Lindemann-Matthies & Marty, 2013), and their ecological value is determined by multiple environmental factors, such as climate, adjacent landscape mosaic/characteristics (Braschler et al., 2020), natural vegetation (Borysiak et al., 2017; Prendergast et al., 2022), and soil composition (Tresch et al., 2018, 2019). Although these intrinsic attributes are mostly beyond owners’ control, the impact of domestic gardens on biodiversity primarily depends on the decisions of garden owners, which dictates design, use, and management (Quistberg et al., 2016). With appropriate management, even small or highly anthropized domestic gardens can support high biodiversity (Donkersley et al., 2023; Griffiths-Lee et al., 2022; Muratet & Fontaine, 2015). On the other hand, numerous anthropogenic stressors (Varga-Szilay et al., 2024), such as the intensity of cultivation (Krištín et al., 2024; van der Veen, 2005), mowing frequency (Lerman et al., 2018) or pesticide use (Tassin de Montaigu & Goulson, 2023), often compromise the ecological value of gardens (Varga-Szilay & Pozsgai, 2022). There is an agreement that the potential ecological value of gardens in conservation correlates positively with the structural diversity of gardens (Majewska & Altizer, 2020), reduced pesticide usage (Tassin de Montaigu & Goulson, 2023), and reduced mowing frequency (Chollet et al., 2018; Lerman et al., 2018; Proske et al., 2022). Ultimately, however, garden management is a series of ’tyranny of small decisions’ (Dewaelheyns et al., 2016) with mixed motivations (Gifford & Nilsson, 2014; Hruška et al., 2021) and external factors such as demographics, household income, neighbours’ expectations, accessibility of information, and education, can influence owners’ behaviour (Goddard et al., 2013; Varga-Szilay et al., 2024). Hence, considering all factors, evaluating the biodiversity values of gardens necessitates considering the garden owners’ management and attitude toward a diverse garden. Yet, the major drivers in management may depend on several societal factors.

In Western Europe, leisure opportunities and recreational activities have long been the primary drivers of gardening, promoting well-being (Beumer, 2018) whereas in Central and Eastern Europe (CEE) gardening for food self-provisioning (FSP) still may dominate (Jehlička et al., 2020, 2021; Smith & Jehlička, 2013). The Central and Eastern European countries share a historical background in FSP, which is thought to be a coping strategy for economic pressures (Alber & Kohler, 2008; Jehlička et al., 2021). However, the previous differences in gardening motivation are gradually beginning to blur with the emergence of new gardening trends (e.g. wildlife-friendly gardening) in both Western and Central- and Eastern Europe (Keshavarz et al., 2016; Poniży et al., 2021) as the demand for ensuring food security and quality in Western countries grows (Church et al., 2015; Glavan et al., 2018; Pourias et al., 2016). At the same time, gardening in CEE countries shifts toward recreational activities (Petzke et al., 2021; Tóth et al., 2018; Trendov, 2018). Yet, the increasing emphasis on nature conservation across all of Europe (Galluzzi et al., 2010; Muller et al., 2010; Vávra et al., 2014), seems to be embraced by CEE countries slower than in their Western counterparts (Hruška et al., 2021; Vávra et al., 2018).

Thus, the divide in the applied gardening practices between East and West remains significant. Indeed, due to the limited access to biodiversity-related information (Coisnon et al., 2019; European Commission, Directorate-General for Communication, 2015), lower willingness to change to environmentally friendly practices, and the extensive use of pesticides (European Commission, Directorate-General for Communication, 2015), CEE appears to be lagging behind in the adoption of biodiversity-friendly practices (for example leave space for wildlife) (European Commission, Directorate-General for Communication, 2015).

As differences between East and West remain, assessing the ecological value of domestic gardens, therefore, requires consideration of location-specific anthropogenic and environmental variables (Varga-Szilay et al., 2024). Whilst there have been numerous studies investigating the role of gardens in maintaining urban biodiversity in Western Europe since the 1990s (Delahay et al., 2023), in CEE these are scarce (but see Varga-Szilay et al., 2024; Varga-Szilay & Pozsgai, 2022).

To address this knowledge gap, here, we seek to investigate how gardens’ parameters, the gardening motivation of garden owners, and their pesticide use habits influence each other in nine Central- and Eastern European countries and explore the differences and similarities between gardens and gardening practices with a potential for maintaining high biodiversity. Since these can fundamentally drive local educational strategies, we particularly aim to pinpoint how geographical differences and sociodemographic parameters best predict domestic gardens’ ecological values, the respondents’ attitudes towards supporting insect pollinators in their gardens, and their pesticide use habits. Hence, insight into the costs and benefits of initiatives for increasing garden diversities is essential for effective planning and management, our imperative is to highlight those areas in CEE where these programs can yield the greatest rewards (e.g. densely populated regions originally hosting high biodiversity).

## 2. Material and methods

### 2.1 Questionnaire design and data collection

An online questionnaire was distributed in nine CEE countries (Croatia, Czechia, Estonia, Hungary, Latvia, Poland, Romania, Slovakia, and Slovenia), all of which were formerly part of the Eastern Block and are currently members of the European Union.

The questionnaire was translated from Hungarian to English and then from English to Croatian, Czech, Estonian, Latvian, Polish, Romanian, Slovak, and Slovenian. The participants could choose the language in which they wanted to complete the questionnaire. It was mandatory to respond to 55 of the total 59 questions, organised into nine sections, which took a maximum of 15 minutes to complete. The questionnaire gathered information about the location and the main characteristics of the garden, the sociodemographic parameters, the garden owners’ motivations, cultivation habits, pesticide usage, environmental awareness, and pollinator-friendly practices. Although participants had to indicate their gender (male, female, other), their highest level of completed education (elementary, middle, postsecondary, postgraduate), and their residency according to the Nomenclature of Territorial Units for Statistics (henceforth NUTS, Eurostat, 2021), all responses were otherwise anonymously recorded. Because of the large number of NUTS-3 regions in Poland, to simplify the questionnaire, here, residencies were recorded at NUTS-2 levels, while the other eight participating countries were recorded at NUTS-3 levels.

The questionnaire was designed using Google Forms and was actively distributed between 26^th^ October 2022 and 18^th^ May 2023, for 90 days in each of the nine participating countries (**Supplementary Material Table 1**). The questionnaire was disseminated through channels such as gardening-oriented websites, and various online social media platforms (for instance Facebook and Instagram) where QR codes and hashtags were used to increase the sharing efficiency. Moreover, through targeted email distributions, outreach efforts extended to professional associations, non-governmental organisations, foundations, and societies dedicated to domestic gardening and environmental conservation.

The participation was voluntary, and a Data Protection and Privacy Statement was available alongside the questionnaire.

### 2.2 Definitions

Not all respondents are, actually, the owners of the gardens they reported on but, for simplicity, we refer to everyone as a ‘garden owner’. For terminological clarity, we used the definitions of ‘pesticides’ and ‘gardening’ as given by Varga-Szilay et al. (2024). Pesticides were defined as ‘all synthetic and non-synthetic products that are used to control pests’, including ‘all commercially available and homemade plant protection products, either those allowed in organic gardening or used in conventional practices’. Gardening was defined as ‘all garden work and all garden care practices, such as the cultivation of flowers, fruits, vegetables, and ornamental plants, mowing, and soil management’ (Varga-Szilay et al., 2024).

### 2.3 Data processing

For the analysis, we only used 43 questions relevant to our aims of the original 59 ones. The original categorical replies of our questionnaire were re-categorised for analytical purposes on a few occasions (see **Supplementary Methods**). NUTS polygons, number of inhabitants, the regional gross domestic product (GDP) in purchasing power standards per inhabitant for each NUTS (PPS, henceforth), and the urban-rural typology for each NUTS-3 region were obtained from ArcGIS Data and Maps (2022a, 2022b) and Eurostat (2022a, 2024), respectively. PPS was only available for NUTS-2 categories. When country capitals with separate NUTS categories were situated within a larger unit, capitals were merged with the largest NUTS surrounding them (CZ, HR, HU, LV, PL, RO). In these cases, for PSS and number of inhabitants, the mean and sum of the merged areas were calculated, respectively. In the case of urban-rural typology data, the originally obtained NUTS-3 categories were merged into NUTS-2 in Poland, and the NUTS polygons of the above mentioned capitals were merged with the surrounding polygons. For this, a weighted mean of the numerised typology index was calculated using the areas of the merged NUTS polygons as weights. The means were rounded to the nearest integer and converted back to categories. Spatial and polygon calculations were conducted using the ‘sf’ (Pebesma, 2018) R package.

### 2.4 Answer-based scoring system

We rated the potential ecological value of the gardens, the garden owners’ motivation for gardening, and pesticide use habits with an answer-based scoring system. For this, we used a total of 32 questions from the original 59 (i.e. questions related to sociodemographic variables were excluded). The garden (GAR) index shows what potential a domestic garden has to build high biodiversity. This index reflects on the structural diversity of the gardens, including garden size, the area of undisturbed patches and plants covering the garden, and the habitat types of adjacent areas, as well as, the disturbance level and the presence of artificial habitats. The calculation of this index was based on 12 questions and consisted of 10 components, which contributed with different weights. The weights were determined by the importance of the components in maintaining high biodiversity (**Supplementary Methods section 1.1**). The respondent (RES) index assesses the garden owners’ knowledge of garden wildlife and their attitudes to maintaining/creating a biodiversity-friendly garden. The RES index was based on 10 questions and consisted of 10 components, including both theoretical (e.g. ‘Do you think…?’/’Can you imagine…?’) and practical (e.g. ‘How do you…?’) questions/question groups, which contributed to the index with different weights (**Supplementary Methods section 1.2**). The pesticide (PES) index shows the degree of pesticide load in gardens by assessing the amount and diversity of pesticides used and the related knowledge of the garden owners. It was based on 10 questions and consisted of 11 components, which were contributed to the index with different weights (**Supplementary Methods section 1.3**). All indices were scaled between 0 (as the lowest) and 100 (as the highest) points. For the full details of the index calculation process, the reader should consult with the **Supplementary Methods**.

### 2.5 Bird index

Since garden diversity depends on the historical biodiversity of a given area (Dobrovodská et al., 2023) but simplified, human-modified landscapes can achieve moderate levels of ecological benefits more rapidly (Haenke et al., 2009), the efficiency of biodiversity-friendly activities can only be measured in the context of local biodiversity indices. Thus, as a proxy for local biodiversity, we collected a full list of bird species recorded between 2010 and 2023 in each NUTS-3 area (except in Poland, see section 2.1) from the Global Biodiversity Information Facility (GBIF.org, 2024). To minimise the observer bias and the effect of rare species, in each NUTS, we only considered species that had at least five recorded occurrences per year from the particular area. We standardised the collated number of species with the area of the NUTS (i.e. divided the species number with the area of NUTS in km squares). Using the same methodology we also gathered a species list for the whole area of interest (i.e. the nine participating countries) and standardised for bird species per km^2^. We then divided the standardised local bird richness with the standardised whole-area bird richness. For collecting bird data we used the ‘rgbif’ (Chamberlain et al., 2024) and for spatial calculations the ‘sf’ (Pebesma, 2018) R packages.

### 2.6 Urban-rural typology

The European urban-rural typology data is a qualitative index which classifies grid cells into rural and non-rural grids and establishes three categories (predominantly rural, intermediate, and predominantly urban) of urbanisation based on the percentage of the population living in rural grid cells (Eurostat, 2024). We employed this index to approximate urbanisation. Since the qualitative nature of this index, we also converted our four indices (bird, GAR, RES, PES) to categorical variables by dividing them at their 0.33 and 0.66 quantiles. When calculating the quantiles, only measured values were included and the theoretical minimum and maximum were not considered. NUTS areas with less than five respondents were excluded.

### 2.6 Statistical analysis

We examined the correlation between the three indices (GAR, RES, PES) with the Spearman correlation test. For calculating the correlation matrix the ‘psych’ (Revelle, 2021) R package was used. We used the Kruskal-Wallis method to test if GAR, RES, and PES indices differ among countries. Pairwise differences were investigated using the pairwise Wilcoxon test with Holm-corrected p-values.

The three indices were used as response variables in three separate Gradient Boosting Machine (GBM) learning processes. Seven sociodemographic variables (gender, age, education level, having children, gardening experience, gardening perception and the average time spent gardening), the PPS, latitude and longitude of the centroids of the NUTS areas were also included in the model as numerical explanatory variables. We randomly divided the datasets into training (70%) and test (30%) sets. After an optimisation and tuning process of the model parameters, we built the GBM model using a Gaussian distribution with 5 levels of interaction depth, with 0.3 shrinkage, 0.80 bag fraction, and a 10 fold cross validation on 28 (for GAR index), 26 (for RES index), and 10 (for PES index) trees. The model fit was evaluated by calculating the R-squared and Root Mean Standard Error (RMSE) values. We used the SHapley Additive exPlanations (SHAP) method to interpret our final GBM models with the ‘shapviz’ (Mayer & Stando, 2023) and ‘kernelshap’ (Mayer et al., 2023) R packages. The SHAP assesses individual variable contributions, accounts for variable interactions, assigns importance values, and facilitates the comparison between all possible variable orders (Lundberg & Lee, 2017). Since GBM predicted the country identity as the most influential variable on all three indices, we examined the relationship between GAR and PES indices (the two non-correlating ones) separately for each country and explored how the other important variables grouped within this data cloud.

For modelling and the visualisation of model results, we used the ‘gbm’ (Greenwell et al., 2022) and ‘caret’ (Kuhn et al., 2023) packages in an R environment (R Core Team, 2021).

Preliminary data clean-up was done in a Python (Python Software Foundation, 2019) environment, with the help of ‘NumPy’ (Harris et al., 2020) and ‘Pandas’ (The pandas development team, 2022) libraries.

## 3. Results

### 3.1 Garden owners’ sociodemographic characteristics

Altogether 5255 garden owners completed the questionnaire from the 9 participating countries (**Supplementary Material Fig. 1, Supplementary Material Table 1**), of which 1146 were males (21.80%), 4094 were females (77.91%), and 15 were non-binary gender (0.29%). The majority of respondents were between 36 and 55 years old (n = 2841, 54.06%). The greatest proportion of garden owners had a postsecondary education (n = 3225, 61.37%), followed by the middle education (n = 1466, 27.90%) and postgraduates (n = 512, 9.74%), while only 52 respondents had elementary school as the highest level of their education (0.99%) (**Supplementary Material Table 2**).

In all participating countries, most of the respondents considered gardening as their favourite hobby (between 21.05% and 58.31%) or a pleasant pastime (between 26.46% and 62.54%) (**Supplementary Material Fig. 2**). In most participating countries, mowing several times a month was under 40% (**Supplementary Material Fig. 3**) but in Estonia and Latvia, the majority of garden owners (65.07% and 63.61%, respectively) mowed the lawn several times a month, and the proportion of those who did so once a month was also high (27.04% and 21.88%). The average pesticide use among the nine countries was 52.60%, ranging from 38.88% in Slovakia to 69.59% in Romania (**Supplementary Material Fig. 4**).

### 3.2 Indices

The mean of the calculated indices was 56.49 (range: 14.97, 91.32), 63.25 (range: 9.25, 96.75), 76.33 (range: 24.87, 100), for GAR, RES, and PES, respectively (**Supplementary Material Table 3**). The GAR index with the RES index showed a moderate, positive correlation (Spearman’s rho = 0.55, p < 0.001). The PES index showed an almost negligible, negative correlation with the RES index (Spearman’s rho = -0.09, p < 0.001). There was no significant correlation between GAR and PES indices (p = 0.487).

There were significant differences between the participating countries in all three indices (GAR, RES, PES) (Kruskal-Wallis chi-squared = 290.85, 671.51, and 235.13, respectively, p < 0.001) (**Supplementary Material Table 4, 5**), and the index values varied broadly among NUTS areas (**Fig. 1**). A total of 197 respondents had all three scores below the first quartile of the indices, with those from Romania and Hungary faring the worst, whilst there were a total of 90 respondents in the fourth quartile of all indices, mostly from Czechia and Slovenia (**Supplementary Material Table 3, 4**).

**Fig. 1:**
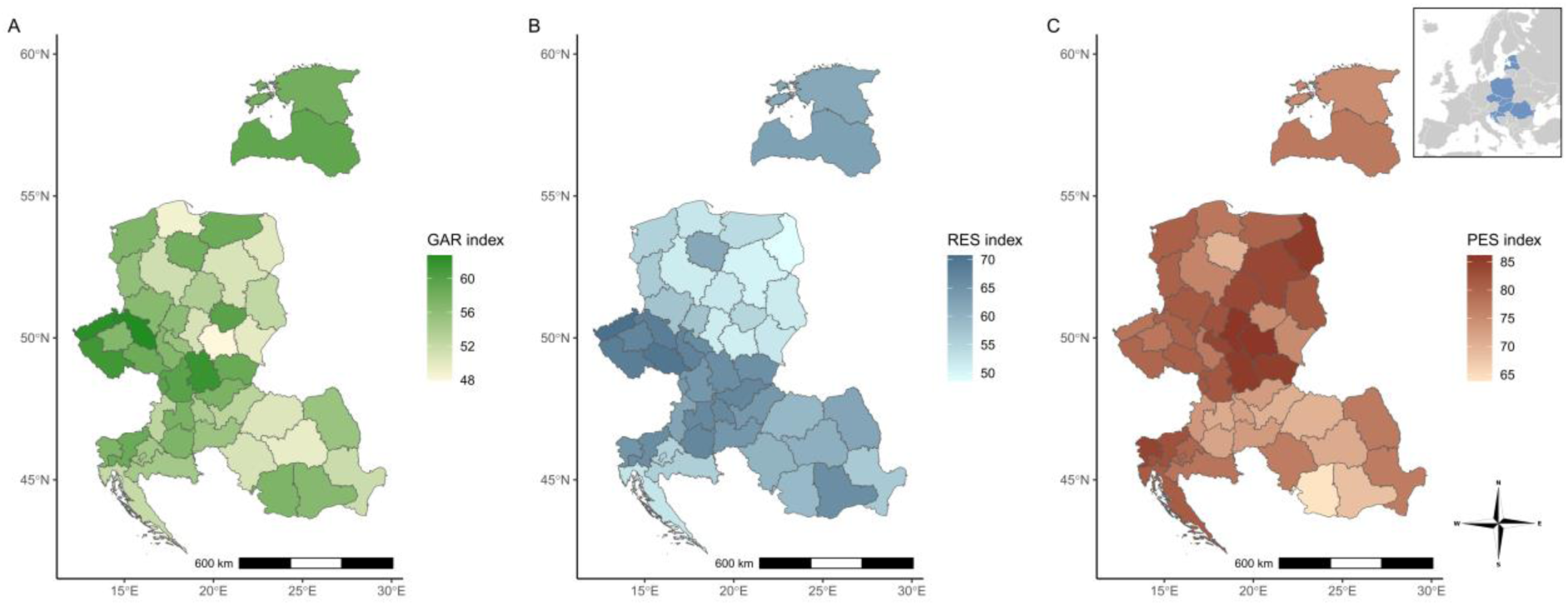
The garden (A), respondents (B) and pesticide (C) indices in the nine participating countries. The colour depth of the maps indicates the mean of the indices calculated for each NUTS-2 area.

### 3.3 GBM outputs

#### 3.3.1 GAR index

The GBM model for the GAR index explained 12.45% of the variance for new observations (RMSE = 11.29), with SHAP values indicating the country variable being the best and the average time that garden owners spent with gardening being the second-best predictor (SHAP value = 2.15, 1.94, respectively) (**Fig. 2A**).

**Fig. 2:**
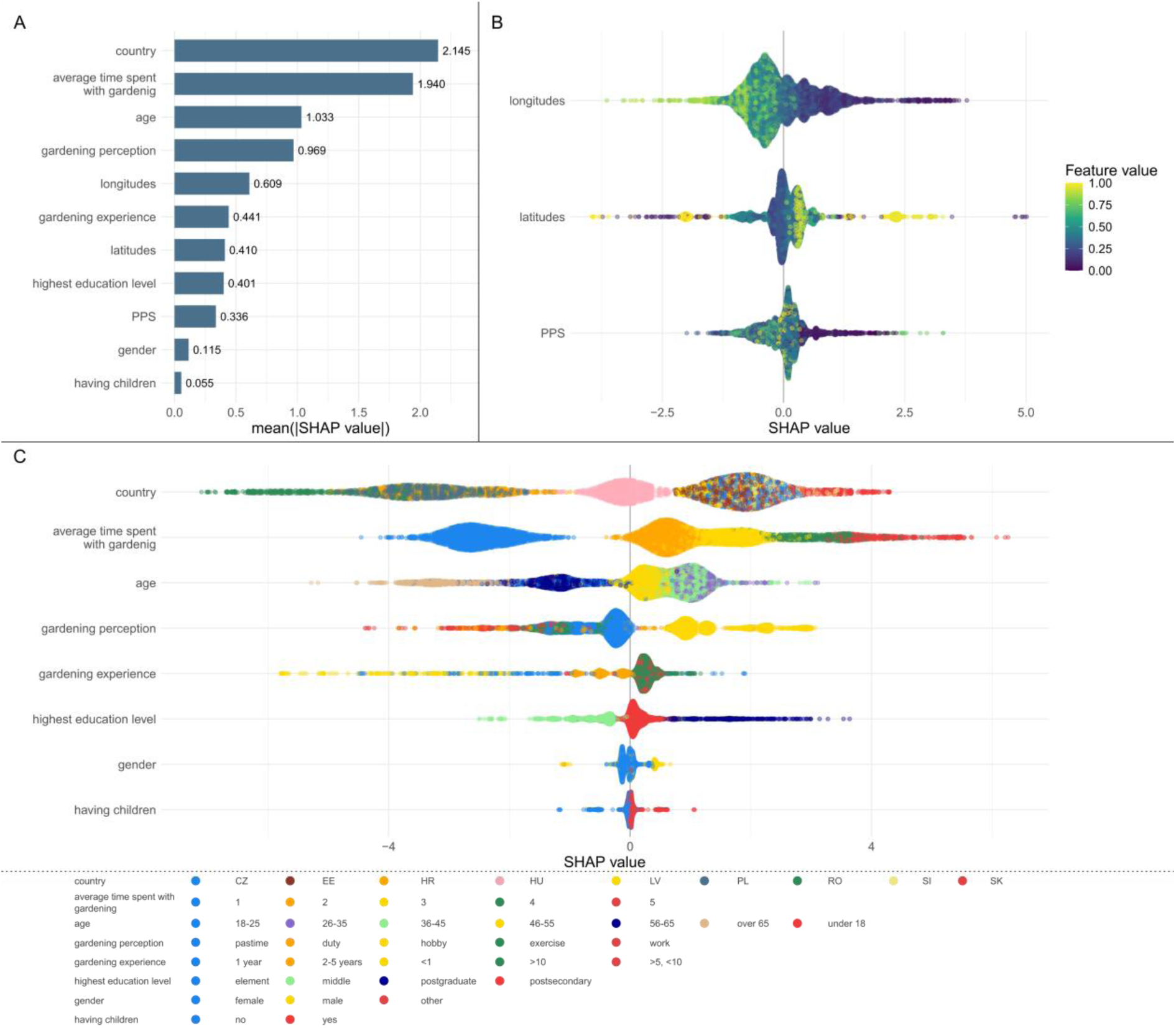
Global SHAP (SHapley Additive exPlanations) summary plots for the GBM model of the GAR index. Variables are ordered by their (A) importance based on average absolute SHAP values. Beeswarm plots of SHAP values of (B) numerical and (C) categorical variables. Each point represents one respondent. For numerical variables, colours indicate variable values, while for categorical variables, they represent distinct category levels. Countries are abbreviated as CZ, EE, HR, HU, LV, PL, RO, SI, and SK, for Czechia, Estonia, Croatia, Hungary, Latvia, Poland, Romania, Slovenia, and Slovakia, respectively. Time spent with gardening is shown in a self-reported scoring system as indicated by the respondents, with 1 representing the lowest (‘one to two hours per day) and 5 representing the highest value (‘even twelve hours per day’).

Countries like Romania, Poland, and Croatia shifted the GAR index toward the lower, while Czechia, Estonia, Latvia, Slovenia, and Slovakia to higher values (**Fig. 2C**). The DIV index increased with increasing time spent with gardening.

The two age categories over 55 shifted the GAR index toward lower values (particularly the age group of over 65), in contrast, the three age categories between 26 and 55 resulted in higher values. Perceiving gardening as a work or a pleasant pastime both lowered the GAR index values while perceiving it as a favourite hobby increased them. Postsecondary and postgraduate education shifted the index to higher values, while middle-level education to lower ones. Albeit it had a low influence in the model, garden owners who had children separated well from those who did not, with those having children shifting the index to higher values. The GAR index values were increased from the west to the east (**Fig. 2B**).

#### 3.3.2 RES index

The GBM model for the RES index explained 24.73% of the variance for new observations (RMSE = 13.13), with SHAP values indicating the country variable being the best and how garden owners perceived gardening being the second-best predictors (SHAP value = 3.80, 2.26, respectively) (**Fig. 3A**). Whilst Hungary and Czechia had the highest SHAP values, and thus the RES index, Poland and Croatia had the lowest ones, with Romania and Latvia also indicating lower values. Only perceiving gardening as a favourite hobby shifted the SHAP values to higher ranges, all other cases lowered them, especially when gardening was perceived as a duty (**Fig. 3C**). SHAP values increased with the time spent with gardening. Older age groups predicted lower RES index values, while the 26-35 age group exhibited higher ones. Gender, education, and having children were less important variables. However, SHAP values separated well with gender, with women presenting at higher values. SHAP values increased from west to east, except for a few very eastern locations presenting the highest values on the right side of the axis (**Fig. 3B**).

**Fig. 3:**
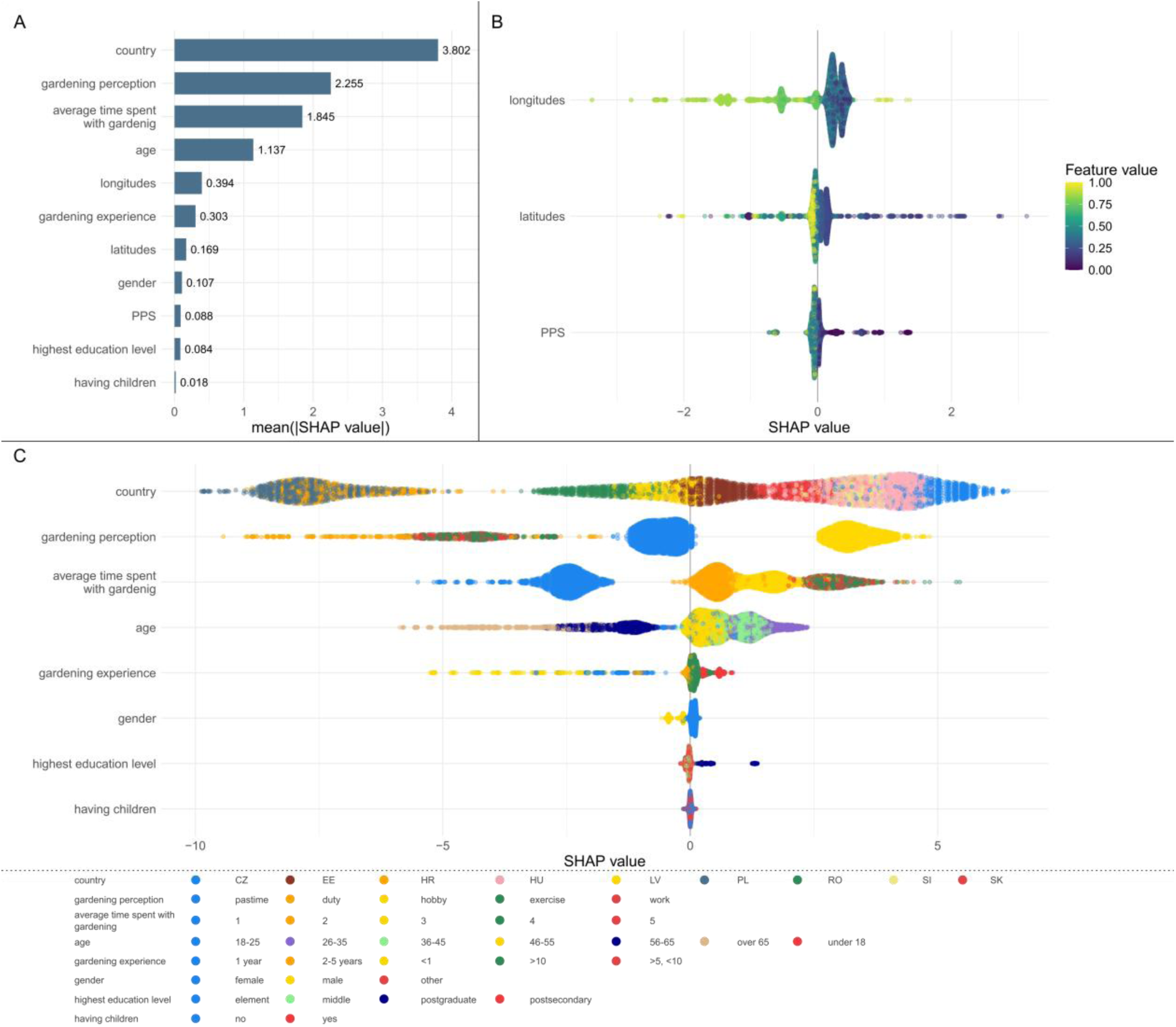
Global SHAP (SHapley Additive exPlanations) summary plots for the GBM model of the RES index. Variables are ordered by their (A) importance based on average absolute SHAP values. Beeswarm plots of SHAP values of numerical (B) and categorical (C) variables. Each point represents one respondent. For numerical variables, colours indicate variable values, while for categorical variables, they represent distinct category levels. Time spent with gardening is shown in a self-reported scoring system as indicated by the respondents, with 1 representing the lowest (‘one to two hours per day) and 5 representing the highest value (‘even twelve hours per day’).

#### 3.3.3 PES index

The GBM model for the PES index explained 0.08% of the variance for new observations (RMSE = 18.19), with SHAP values indicating the country variable being the best and the average time that garden owners spent with gardening being the second best predictors (SHAP value = 3.20, 1.56, respectively) (**Fig. 4A**). Romania, Estonia, and Hungary had the lowest SHAP values, whilst Poland, Slovakia, and Slovenia showed the highest ones (**Fig. 4C**). An average of 1–2 hours of gardening per day predicted a higher PES index, whilst all other times spent with gardening lowered it, with the lowest values associated with those who spent up to 12 hours a day with gardening. More than 10 years of gardening experience shifted the index to lower values. Perceiving gardening as a pleasant pastime shifted SHAP to higher values, whereas in most other cases, the values decreased equally when gardening was perceived as a duty and a favourite hobby. Women shifted SHAP to higher values, however, gender, age, having children, and education level were not deemed to be important variables. Having children separated positive and negative SHAP values, with those respondents who had children associated with higher values. SHAP values increased both from south to north and east to west (**Fig. 4B**).

**Fig. 4:**
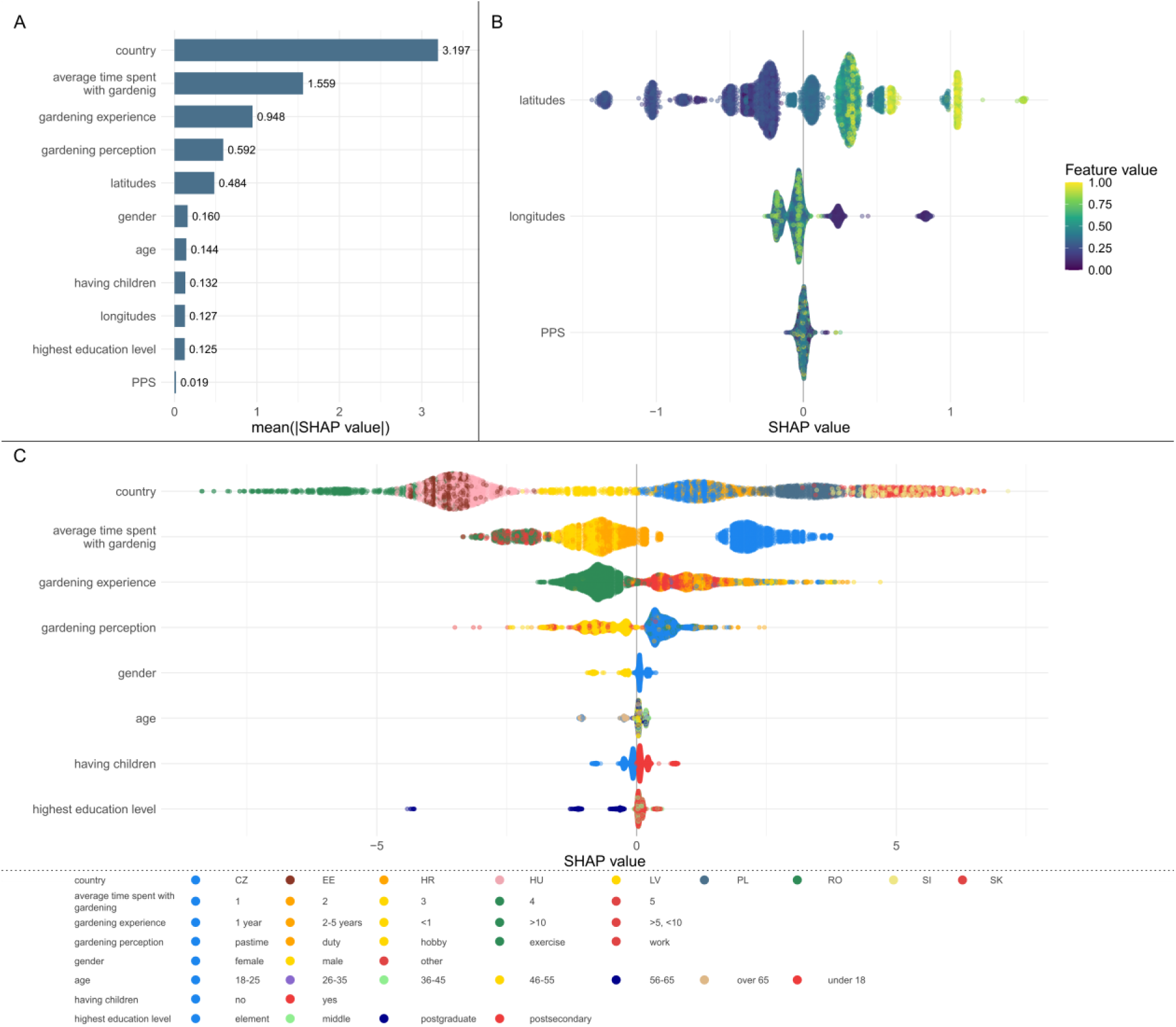
Global SHAP (SHapley Additive exPlanations) summary plots for the GBM model of the PES index. Variables are ordered by their (A) importance based on average absolute SHAP values. Beeswarm plots of SHAP values of numerical (B) and categorical (C) variables. Each point represents one respondent. For numerical variables, colours indicate variable values, while for categorical variables, they represent distinct category levels. Time spent with gardening is shown in a self-reported scoring system as indicated by the respondents, with 1 representing the lowest (‘one to two hours per day) and 5 representing the highest value (‘even twelve hours per day’).

#### 3.3.4 The GAR index as a function of the PES index

In our investigation into the influence of country, GAR index, and PES index on the grouping of gardening experience lengths, we found that respondents who had been gardening for more than 2 years were characterised by a greater potential for building high biodiversity and more intensive pesticide use. In Estonia, Latvia and Slovakia, those who had been gardening for less than a year grouped around low pesticide input (high PES index values). On the other hand, in Slovenia, among those who had been gardening for a year or less, there was no one who did not use pesticides at all. In the case of Hungary and Estonia, gardeners who have been gardening for less than one year scored lower in the GAR index but were also characterised by lower PES index values. (**Supplementary Material Fig. 5**).

In Hungary, Poland, Romania, and Slovenia for those garden owners who considered gardening as their favourite hobby the GAR index was greater than in other countries and they also scored lower in the PES index (**Supplementary Material Fig. 6**).

In all participating countries, except Slovakia, the GAR index was the lowest and the PES index the highest among garden owners who spent less than 1-2 hours with gardening (**Supplementary Material Fig. 7**).

In all countries but Croatia older age negatively influenced the GAR index. Indeed in Romania, garden owners over the age of 65 fared poorly with the GAR index and all of them used pesticides. In contrast in Croatia, this age group had the highest GAR index values (**Supplementary Material Fig. 8**).

### 3.4 Areas with a potential for improvement

When areas within urbanisation levels were cross-referenced with the categorised three indices, different effects of urbanisation levels of the three indices were revealed (**Fig. 5**). Half of the predominantly urban NUTS areas (n = 3) were categorised as low GAR values, and half of the NUTS areas typologised (n = 26) as intermediate reached a medium level on the GAR index. The RES categories were relatively evenly spread both among intermediate and predominantly rural areas but most predominantly urban regions (n = 4) were categorised as medium RES (**Table 1**). The distribution of PES categories was uniform among predominantly urban and predominantly rural areas, whilst in intermediately urbanised regions intermediate PES category was the most common (n = 22), followed by the high and low categories (n = 17 and 13, respectively) (**Table 1**, **Fig. 5**).

**Fig. 5:**
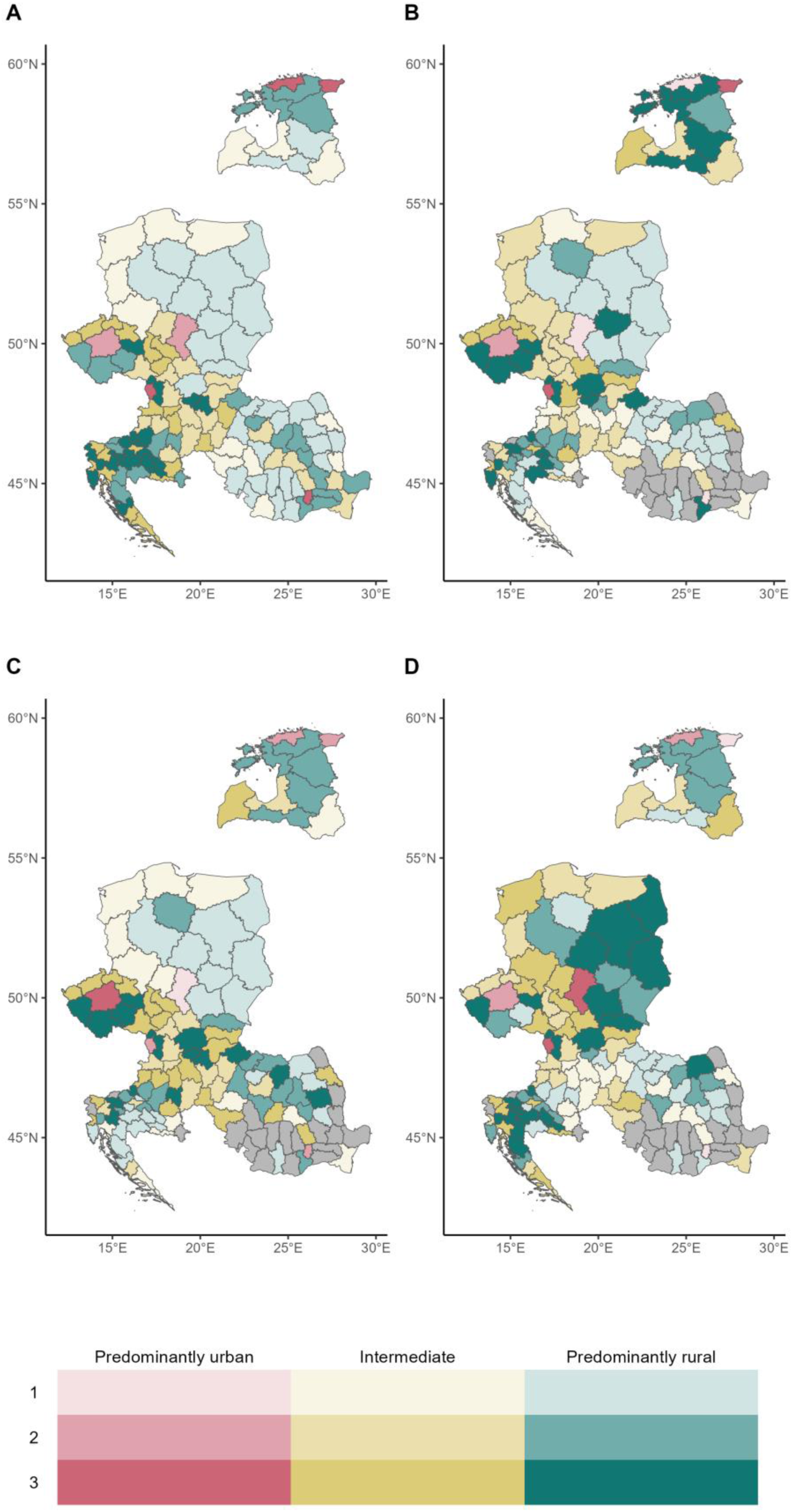
The relationship of the bird (A), garden (B), respondent (C), and pesticide (D) indices with the urban-rural typology index based on NUTS-3 areas **(except for Poland, in** which NUTS-2 were used). The colours represent the levels of urban-rural typology, as follows: red – predominantly urban, yellow – intermediate, green – predominantly rural. The transparency of the colour indicates the categorised index value (bird/GAR/RES/PES) on a three-level scale (divided into categorised at the 0.33 and 0.66 quantiles). The grey colour indicates NUTS areas from which we had less than five respondents (n = 20).

**Table 1:**
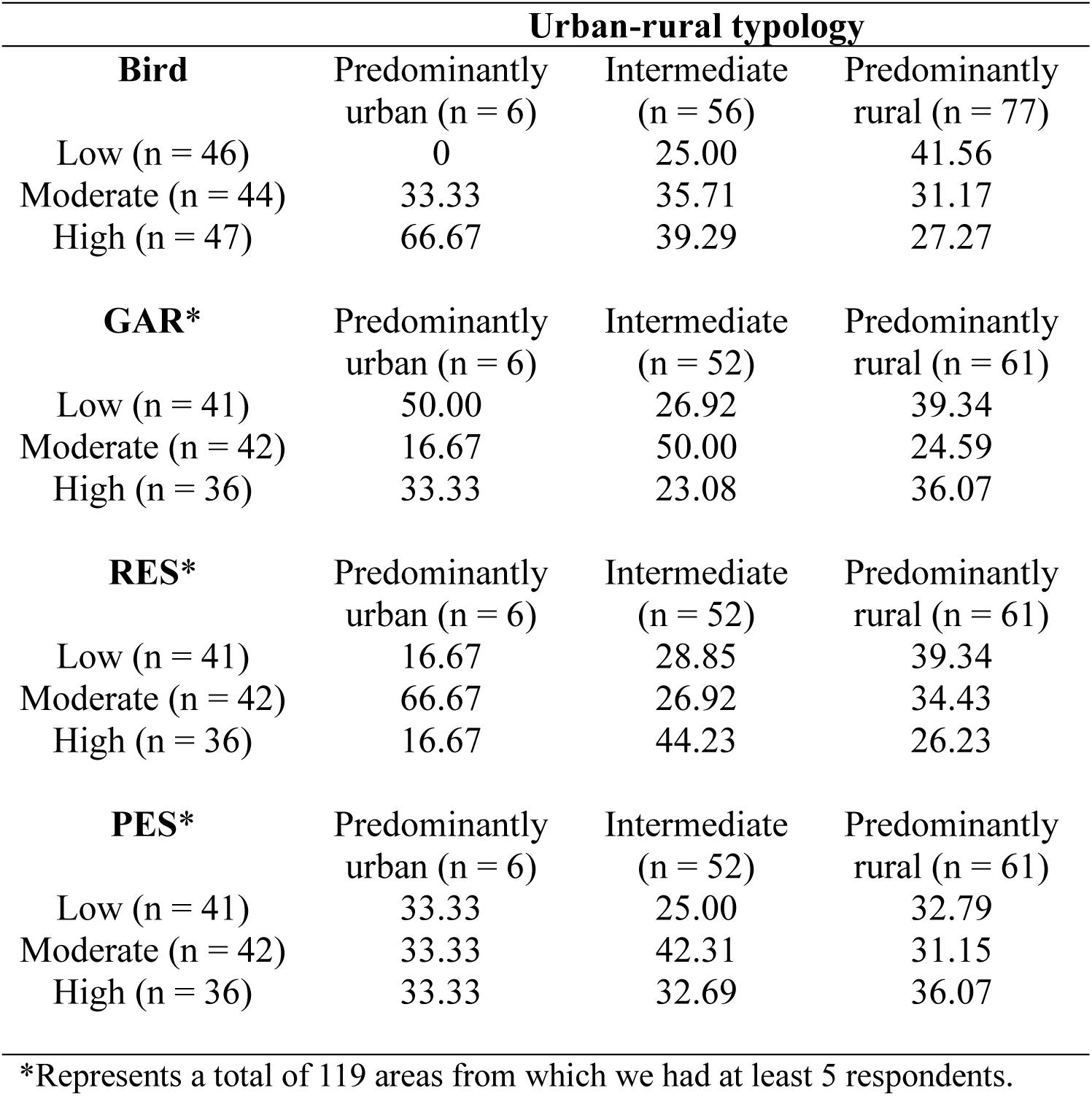
Contingency tables of urban-rural typology categories and categorised index values of bird, GAR, RES, and PES indices, as expressed by the number of NUTS areas. Values are shown as percentages of the columns.

|  | **Urban-rural typology** |  |  |
| --- | --- | --- | --- |
| **Bird** | Predominantly urban (n = 6) | Intermediate (n = 56) | Predominantly rural (n = 77) |
| Low (n = 46) | 0 | 25.00 | 41.56 |
| Moderate (n = 44) | 33.33 | 35.71 | 31.17 |
| High (n = 47) | 66.67 | 39.29 | 27.27 |
| **GAR\*** | Predominantly urban (n = 6) | Intermediate (n = 52) | Predominantly rural (n = 61) |
| Low (n = 41) | 50.00 | 26.92 | 39.34 |
| Moderate (n = 42) | 16.67 | 50.00 | 24.59 |
| High (n = 36) | 33.33 | 23.08 | 36.07 |
| **RES\*** | Predominantly urban (n = 6) | Intermediate (n = 52) | Predominantly rural (n = 61) |
| Low (n = 41) | 16.67 | 28.85 | 39.34 |
| Moderate (n = 42) | 66.67 | 26.92 | 34.43 |
| High (n = 36) | 16.67 | 44.23 | 26.23 |
| **PES\*** | Predominantly urban (n = 6) | Intermediate (n = 52) | Predominantly rural (n = 61) |
| Low (n = 41) | 33.33 | 25.00 | 32.79 |
| Moderate (n = 42) | 33.33 | 42.31 | 31.15 |
| High (n = 36) | 33.33 | 32.69 | 36.07 |

\*Represents a total of 119 areas from which we had at least 5 respondents.

Of the NUTS areas typologised as predominantly urban, Czechia, Hungary, Slovakia and Slovenia had those with the highest scores of all calculated indices (bird, GAR, RES, and PES) (e.g. the Plzeň region in Czechia reached the best categories in all variables but the bird index), while Romania had that with the lowest (the Vrancea county in Romania fall into the worst categories in all variables). All predominantly urbanised areas were categorised relatively high in all indices with Slovakia being the best (Bratislava region) and Poland and Romania being the worst (Silesian Voivodeship, București-Ilfov region).

## 4. Discussion

Domestic gardens offer significant opportunities for advancing global biodiversity and sustainability targets. Our work investigated the intertwined relationship between sociodemographic parameters, sustainable garden management, and biodiversity conservation from a highly understudied area of the European Union and highlights that there are significant differences among countries, and even within countries, regardless of the scale, in the examined parameters.

Indeed, the participating countries differed in all three examined indices, with having the least differences in GAR and the most in PES, and these differences were supported by both the pairwise comparisons between countries and the GMB model. There were countries experiencing either equally low or high values on all indices, such as Romania faring poorly in all indices, whereas Slovakia showing overly high values. Sometimes indices implied counteracting effects, such as a high GAR index, suggesting a great prospect for maintaining high biodiversity in domestic gardens, occurred together with high pesticide use in Hungary. This is mostly in line with Coisnon et al. (2019) who found similar trade-offs in variables influencing sustainable gardening practices such as Hungary scoring the worst in leaving spaces for wildlife and avoiding the use of pesticides, yet having been categorised as the best in selecting plants that provide food for birds and pollinators and avoiding the introduction of potentially invasive plants. Other countries such as Romania generally ranked poorly in both our study and known from the literature (Coisnon et al., 2019). All three indices varied widely, however, indicating the multidimensional nature of factors driving sustainable gardening.

The extensive application of pesticides beyond the agricultural sector, including domestic gardens, and the inter-country disparities in pesticide usage within Europe are critical issues for biodiversity-friendly gardens (Poniży et al., 2021; Varga-Szilay et al., 2024; Varga-Szilay & Pozsgai, 2022). Our study found substantial differences between countries in this regard, with 20–30% of respondents in Hungary and Romania thinking that the use of pesticides is either important or crucially important for their gardening, a claim with which only 2% of those from Slovakia agreed (**Supplementary Material Fig. 9**.).

There is a disparity though between pesticide use and the other two indices we measured. Although nearly 70% of respondents from Romania considered the widespread use of pesticides as one of the most threatening factors to pollinators, their average pesticide use was still the least favourable in our study. Indeed, what gardeners perceive as a biodiversity-friendly garden may not necessarily align with its real biodiversity potential, suggesting that self-reporting surveys may be misleading in this term, unless gardens are examined from multiple aspects, including sociodemographic or cultural factors.

Indeed, in line with Larson et al. (2022), those who perceived gardening as a hobby were more inclined to change their gardens to be attractive and beneficial to biodiversity, and we also discovered that garden owners who dedicate more time spent with gardening tend to maintain gardens with higher biodiversity potential. Our results corroborate with the findings of both Philpott et al. (2020) and Lin et al. (2017) who indicated that time spent in domestic gardens positively correlates with plant species richness, higher dissimilarity in crop composition, and high levels of nature-relatedness. Moreover, our findings agree with those of Geppert et al. (2024) who reported that the time spent outdoors increased the willingness to pro-pollinator actions. In our study, however, both when gardening was associated with a hobby and more time was allocated for gardening were characterised by increased pesticide use. Therefore, there is a higher possibility for these gardeners to create an ecological trap in these gardens for visiting organisms (especially insects) (Varga-Szilay & Pozsgai, 2022). There is, however, a noticeable contrast in behaviour based on gender and parental status (**Fig. 4C**) where women and respondents with children improve the outcome of the PES index, whilst this separation was unclear with the other two indices.

Although the country identity proved to be the most important variable in predicting the potential for maintaining high garden biodiversity, the respondents’ attitudes towards supporting biodiversity (especially pollinators) in their gardens, pesticide use, owners’ perception, the long-term gardening experience and daily management also proved to be important factors. Furthermore, nature conservation was the prime aspect that influenced gardening habits in four participating countries, albeit in three of these this did not mean a lower pesticide use, and only Croatia and Poland were the ones where more than 80% of the respondents felt that nature conservation does not affect their gardening habits at all (**Supplementary Material Fig. 10**). Increasing age (especially that over 65) negatively influenced both the potential of gardens for maintaining high biodiversity and the owners’ attitudes toward maintaining biodiversity-friendly gardens, which suggests the elderly are more inclined to follow and uphold longstanding conventional practices, regardless of whether or not they are beneficial for ecological sustainability. Whether these decisions are driven either by the difficulties the elderly may have with manoeuvring among the continuous flow of complex ecological issues which were not highlighted during most of their lives, the scarcity of the available information from the more traditional information channels they use, or some alternative explanations is yet to be clarified. Nonetheless, the age-independent pesticide use we observed in the GBM model suggests that even when information is broadly available behavioural change is not granted.

However, gardening practices and their associated sociocultural and economic variables exhibited significant diversity among garden owners and the final outcome for biodiversity-friendliness appeared to be a series of multifactor decisions depending on numerous background drivers (Gifford & Nilsson, 2014). For instance, despite several studies highlighting a substantially positive effect of GDP per capita and household income on sustainable gardening practices and pollinator abundance in domestic gardens (e.g. Baldock et al., 2019; Coisnon et al., 2019), in our GBM model, PPS did not prove to be a significant explanatory variable for any of our indices.

Despite the existing differences among the NUTS regions and countries in all three indices, we did not find strong geographical correlations; neither latitude nor longitude had a high explanatory power in our GBM models. Nonetheless, longitude had a higher influence on the GAR index than latitude, which might not be as expected since the north-south gradient exhibits steeper changes in climate and habitats, than the one from east to west. This greater importance of the east-west axis could be explained by the higher impact of Western European countries (and western culture) on how more westerly regions in CEE perceived biodiversity issues after political and societal changes in the 1990s. Conversely, the lower influence of longitude and latitude on the RES and PES indices could be attributed to the continued strong connection to traditional practices across the studied countries.

Our results show that merely examining the structural diversity of private gardens, sociodemographic parameters, garden owners’ motivation for gardening, or biodiversity-negative activities (e.g. pesticide use) individually may be insufficient for obtaining a reliable proxy of whether domestic gardens could potentially maintain/build high diversity in human-altered areas. Moreover, the lack of a clear geographical pattern and within-country diversity in all indices suggests that regionally tailored rather than countrywide strategies are needed to identify pathways toward sustainable green infrastructure and to improve the natural value of domestic gardens. Since these strategies must be underpinned by interdisciplinary study efforts, our study can offer fundamental insight into how they can be optimised and where the greatest rewards can be gained. Indeed, despite the dominance of moderate values of our indices on the map, local strategies can still be developed based on the preliminary classification.

Areas with generally **low urbanisation** and also **low biodiversity** are likely to be agricultural- and farmlands in which gardens with high biodiversity can act as habitat islands, ecological corridors, or stepping stones which facilitate the movement of wildlife through unsuitable agricultural areas. In these areas, improving the structural complexity of the gardens and reducing pesticide use could be the most beneficial for biodiversity. Disseminating knowledge about ecological-friendly practices and sustainable gardening habits would also be key. Indeed, the predominantly rural areas in northeast Poland, where both bird index and urbanisation are low, also suffer from low RES. However, information exchange here may need to rely more on interpersonal relationships or traditional media than online channels (Troumbis, 2021) and overall biodiversity is likely to be more impacted by agricultural rather than garden practices. Attempting to improve biodiversity value in areas with generally **low urbanisation** levels and **high biodiversity** probably has the least merit, yet reducing pesticide use can prohibit degradation and may form garden-level biodiversity islands that increase landscape complexity. Moreover, maintaining and even improving garden diversity in these areas, such as predominantly rural areas in the northern part of Croatia and Slovenia, can be particularly important as good examples for environmental education and maintaining source populations of wildlife for adjacent but less favourable areas.

Thus, at low urbanisation levels improving aspects of the RES would be the priority, along with a substantial decrease of pesticide use. With increasing biodiversity, gardens will benefit from improved conservation biological control (Lacey et al., 2015; Landis et al., 2000; Quesada- Moraga et al., 2022), which may further decrease pesticide use.

Biodiversity-friendly activities can achieve fast and great ecological benefits in areas where **urbanisation** is **high** and **biodiversity** is **moderate or low**. Although their biodiversity potential will remain limited, prioritising increasing gardens’ structural and plant diversity will enable them to serve as ecological corridors or stepping stones for most wildlife to navigate around highly urbanised areas. Although biodiversity will unlikely increase substantially, the high visibility of biodiversity in densely populated areas has an outstanding educational value. Even though few CEE regions fulfil these criteria (e.g. the central part of Czechia), most Western European areas predominantly classified as urban are likely to fall into this category. Although **high urbanisation** and **high biodiversity** rarely co-occur, some areas in CEE, such as Northern Estonia, exhibit such a phenomenon. This offers an opportunity for the dissemination of activities tied to increasing the RES index (e.g. through campaigns and species identification trainings), as they can quickly reach many individuals and engage them in enhancing garden biodiversity. Additionally, these efforts provide a substantial amount of data for biodiversity monitoring, resulting in additional benefits.

Thus, at high urbanisation levels, increasing gardens’ structural diversity and other aspects beneficial for maintaining high biodiversity are key but improving garden owners’ attitudes towards biodiversity-friendly gardening and reconnecting them to nature are also important. Since the level of urbanisation across Europe is projected to reach 83.7% by 2025 (United Nations, 2018), this improvement in attitude is necessary to exploit the potential of domestic gardens to mitigate large-scale biodiversity loss. However, due to the ubiquitous use of pesticides in CEE, improving the PES index would be equally necessary in every area, both urban and rural.

Adequate planning, at least, at the regional scale, with the active involvement of garden owners, is crucial for increasing the biodiversity value of domestic gardens at a larger scale. For this, future nature conservation planning strategies and studies should acknowledge the variability of domestic garden owners’ attitudes and gardening practices, as well as the domestic gardens’ potential for biodiversity conservation (Cameron et al., 2012).

### 4.1 Study limits

Although in our work we collected over 5255 responses, this only represents a small fraction of the inhabitants and gardeners of the focal area of our study. Moreover, the dissemination channels we used do not allow random respondent selection and our selection process is most likely to be biased towards those who had regular access to the Internet, whilst those living in infrastructurally less developed parts of the countries had a lower chance to learn about our call. Indeed, parts of Poland and the Southern part of Romania are highly underrepresented. Yet, several countries like Estonia and Hungary, have a high coverage which likely portrays faithfully the common gardening behaviours in those regions.

In an effort to engage a broad audience in several languages, we often compromised completeness for simplicity and brevity. Whilst we believe this did not affect the interpretability of our questionnaire, it definitely sets limits on the use of scientific terminology and how detailed information we could gain with some questions.

Although our questionnaire aimed for the response of only one person (the garden owner) for each domestic garden, these gardens are probably cared for by more than one person (e.g. family members), and thus, they are not only influenced by one person’s motivation but by severals’.

### 4.2 Future perspectives

Involving more participants from the surveyed CEE countries, standardising the questions and expanding the area of interest to all European countries, as well as repeating the same survey within a reasonable time frame would all be highly advantageous. Especially investigating the distinction between the gardening habits in Western and Central-Eastern European regions, as well as those from the Balkan, such as Serbia and Bulgaria, holds particular potential. While addressing privacy concerns may present challenges, extending our study with measures of biodiversity and landscape configuration from each garden could introduce extra layers of complexity to our exploration.

Furthermore, a direct link between domestic garden diversity, and the potential of thereof, and biodiversity conservation should be established and implemented into urban green space management.

## 5. Conclusions

While gardens may not replace undisturbed natural habitats, it is imperative to reconsider maintaining high levels of urban and rural biodiversity as valuable complements for increasing the area of semi-natural habitat patches and creating a network of ecological corridors and stepping stones (Beninde et al., 2015). In our study, we, however, show that the potential of domestic gardens for maintaining high biodiversity varies widely among CEE regions, and along with it does their conservation value. Therefore, to maximise this value improving several aspects of CEE gardening practices in concert is necessary. The variability both between and within participating countries, underscores the need for approaches tailored to regional circumstances rather than unified regulations across European countries.

Despite some differences observed between north to south, the idea of promoting biodiversity and nature conservation seems to be spreading from the west towards the east and therefore should continue with additional efforts towards the eastern countries. Besides promoting the importance of biodiversity, some concrete measures in agricultural practice and activities should be promoted as well and offered to gardeners to speed up in situ changes, as gardeners still seem to incline more to traditional approaches. The nature-friendly practice seems to be supported more by the higher education levels and by the factor of age, or parenting, those facts can be used to target the promotion of gardening in nature conservation in this part of Europe. On the other hand, to ensure that domestic gardens truly contribute positively to sustainable urban environments, the multifaceted issue of pesticide usage that permeates various aspects of gardening, from food production in allotments to the care of ornamental plants and flea treatments for pets, must be uniformly addressed across all European Union countries. Similarly, information for providing a comprehensive understanding among the public regarding the potential environmental and human health impacts of pesticides should also be made broadly available.

Biodiversity-friendly gardens also improve human health and well-being (Samus et al., 2022) and provide urban residents with the opportunity to reconnect to the natural world (Cameron et al., 2012). Thus, while increasing the diversity of domestic gardens alone may not solve the biodiversity crisis, by creating ecological corridors it can contribute to conservation, foster a shift in human attitudes towards preserving natural areas, and enhance the ecological quality of urban environments. To fully exploit all benefits of domestic gardens and to increase people’s knowledge concerning the environment, population-wide, regular, and high-quality education and community programs which, shape people’s attitudes, willingness to take action, and interest, are needed (Knapp et al., 2021; Shwartz et al., 2012; Sturm et al., 2021; Tikka et al., 2000). Similar programs have been running for decades in Western European countries but they are still scarce in CEE (European Commission, Directorate-General for Communication, 2015; Jehlička & Jacobsson, 2021). Yet, the need for improving the ecological quality of domestic gardens is not unique to Western Europe, where it gets the most emphasis, but is also present in CEE. Hence, local strategies for environmental education emphasising the importance of biodiversity-friendly gardening should be developed for CEE countries as well. These, however, should be tailored to local needs and provide access to biodiversity-related information through a broad variety of channels to reach all strata of society.

Besides serving as a proxy to indicate gardens’ environmental quality, our study also can help in accessing these gardens’ potential in designing eco-networks of biodiversity-friendly gardens. It also can act as a tool for facilitating the decision for optimal strategies to maximise the environmental benefits of domestic gardens.

## CRediT authorship contribution statement

− **Zsófia Varga-Szilay:** Conceptualization; Data curation; Formal analysis; Investigation; Methodology; Project administration; Resources; Software; Validation; Visualization; Roles/Writing - original draft; and Writing - review & editing

− **Arvīds Barševskis:** Data curation; Investigation; Methodology; Project administration; and Writing - review & editing

− **Klára Benedek:** Data curation; Investigation; Methodology; Project administration; and Writing - review & editing

− **Danilo Bevk:** Data curation; Investigation; Methodology; Project administration; and Writing - review & editing

− **Agata Jojczyk:** Data curation; Investigation; Methodology; Project administration; and Writing - review & editing

− **Anton Krištín:** Data curation; Investigation; Methodology; Project administration; and Writing - review & editing

− **Jana Růžičková:** Data curation; Investigation; Methodology; Project administration; and Writing - review & editing

− **Lucija Šerić Jelaska:** Data curation; Investigation; Methodology; Project administration; and Writing - review & editing

− **Eve Veromann:** Data curation; Investigation; Methodology; Project administration; and Writing - review & editing

− **Silva Vilumets:** Data curation; Investigation; Methodology; Project administration; and Writing - review & editing

− **Kinga Gabriela Fetykó:** Conceptualization; Data curation; Investigation; Methodology; Project administration; Validation; and Writing - review & editing

− **Gergely Szövényi:** Conceptualization; Data curation; Investigation; Methodology; Project administration; Supervision; Validation; and Writing - review & editing

− **Gábor Pozsgai:** Conceptualization; Data curation; Formal analysis; Investigation; Methodology; Project administration; Resources; Software; Supervision; Validation; Visualization; Roles/Writing - original draft; and Writing - review & editing

## Declarations

### Data and code availability

The data and the underlying computer code is available in the GitHub repository

https://github.com/zsvargaszilay/domestic_gardens_in_CEE

## Acknowledgements

We are grateful to Geoff Gurr and Stephen Venn for their comments on an earlier version of the manuscript.

