## Supplementary Material for "Improving biodiversity in Central and Eastern European domestic gardens needs regionally scaled strategies"

**This file includes:**

Supplementary Material Figure: 1-10

Supplementary Material Table: 1-7

**Abbreviations**

CEE: Central and Eastern Europe

CZ, EE, HR, HU, LV, PL, RO, SI, SK: Czechia, Estonia, Croatia, Hungary, Latvia, Poland, Romania, Slovenia, Slovakia

NUTS: The Nomenclature of Territorial Units for Statistics

PPS: purchasing power standards per inhabitant

GAR index: garden index

RES index: respondent index

PES index: pesticide index

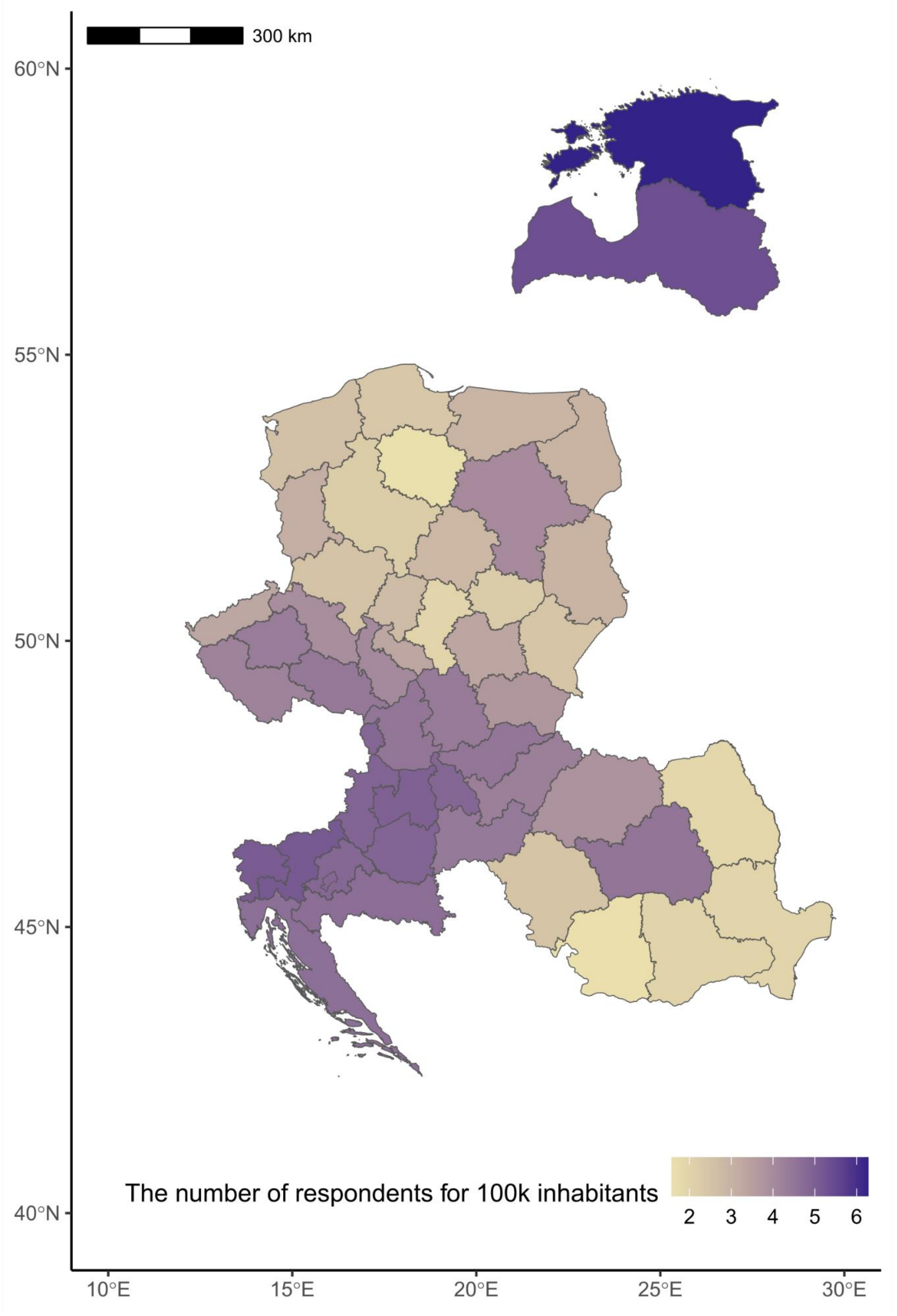


**Supplementary Material Fig. 1:** Number of respondents in the nine participating countries by NUTS-2 areas. The colour depth of the maps indicates the number of respondents by 100 000 inhabitants in each NUTS-2 area. The values are on a logarithmic scale.

**Supplementary Material Table 1:** The first and last days of the data collection period (three months) in the nine participating countries and the number of respondents.

| Country | | **First day* of the data collection** | **Last day of the data collection** | Number of respondents |
|---|---|---|---|---|
| Czechia | CZ | 18/01/2023 | 21/04/2023 | 648 |
| Estonia | EE | 22/01/2023 | 25/04/2023 | 710 |
| Croatia | HR | 27/01/2023 | 30/04/2023 | 462 |
| Hungary | HU | 26/10/2022 | 27/01/2023 | 1088 |
| Latvia | LV | 23/01/2023 | 26/04/2023 | 393 |
| Poland | PL | 14/02/2023 | 18/05/2023 | 746 |
| Romania | RO | 16/01/2023 | 19/04/2023 | 434 |
| Slovenia | SL | 18/01/2023 | 21/04/2023 | 347 |
| Slovakia | SK | 25/01/2023 | 28/04/2023 | 427 |
| **SUM** | | | | **5255** |

*The day on which the first questionnaire was completed.

### Garden owners' sociodemographic characteristics

**Supplementary Material Table 2:** Sociodemographic characteristics of the study population (n = 5 255).

| | | CZ | | EE | | HR | | HU | | LV | | PL | | RO | | SI | | SK | |
|---|---|---|---|---|---|---|---|---|---|---|---|---|---|---|---|---|---|---|---|
| | Levels | n | % | n | % | n | % | n | % | n | % | n | % | n | % | n | % | n | % |
| Gender | Male | 125 | 19.29 | 45 | 6.34 | 90 | 19.48 | 296 | 27.21 | 44 | 11.2 | 212 | 28.42 | 142 | 32.72 | 57 | 16.43 | 135 | 31.62 |
| | Female | 518 | 79.94 | 663 | 93.38 | 372 | 80.52 | 791 | 72.7 | 348 | 88.55 | 531 | 71.18 | 291 | 67.05 | 290 | 83.57 | 290 | 67.92 |
| | Other | 5 | 0.77 | 2 | 0.28 | 0 | 0 | 1 | 0.09 | 1 | 0.25 | 3 | 0.4 | 1 | 0.23 | 0 | 0 | 2 | 0.47 |
| Age | Under 18 | 0 | 0 | 0 | 0 | 1 | 0.22 | 4 | 0.37 | 1 | 0.25 | 2 | 0.27 | 1 | 0.23 | 1 | 0.29 | 0 | 0 |
| | 18-25 | 33 | 5.09 | 4 | 0.56 | 16 | 3.46 | 37 | 3.4 | 6 | 1.53 | 104 | 13.94 | 28 | 6.45 | 10 | 2.88 | 10 | 2.34 |
| | 26-35 | 177 | 27.31 | 89 | 12.54 | 68 | 14.72 | 169 | 15.53 | 43 | 10.94 | 91 | 12.2 | 73 | 16.82 | 49 | 14.12 | 58 | 13.58 |
| | 36-45 | 210 | 32.41 | 180 | 25.35 | 123 | 26.62 | 290 | 26.65 | 63 | 16.03 | 211 | 28.28 | 159 | 36.64 | 94 | 27.09 | 132 | 30.91 |
| | 46-55 | 127 | 19.6 | 216 | 30.42 | 146 | 31.6 | 288 | 26.47 | 128 | 32.57 | 183 | 24.53 | 93 | 21.43 | 88 | 25.36 | 110 | 25.76 |
| | 56-65 | 58 | 8.95 | 166 | 23.38 | 81 | 17.53 | 152 | 13.97 | 119 | 30.28 | 82 | 10.99 | 48 | 11.06 | 65 | 18.73 | 74 | 17.33 |
| | Over 65 | 43 | 6.64 | 55 | 7.75 | 27 | 5.84 | 148 | 13.6 | 33 | 8.4 | 73 | 9.79 | 32 | 7.37 | 40 | 11.53 | 43 | 10.07 |
| Education level | Elementary | 7 | 1.08 | 11 | 1.55 | 0 | 0 | 10 | 0.92 | 5 | 1.27 | 9 | 1.21 | 2 | 0.46 | 4 | 1.15 | 4 | 0.94 |
| | Middle | 229 | 35.34 | 177 | 24.93 | 166 | 35.93 | 294 | 27.02 | 77 | 19.59 | 178 | 23.86 | 120 | 27.65 | 92 | 26.51 | 133 | 31.15 |
| | Postsecondary | 309 | 47.69 | 459 | 64.65 | 241 | 52.16 | 714 | 65.62 | 275 | 69.97 | 507 | 67.96 | 239 | 55.07 | 229 | 65.99 | 252 | 59.02 |
| | Postgraduate | 103 | 15.9 | 63 | 8.87 | 55 | 11.9 | 70 | 6.43 | 36 | 9.16 | 52 | 6.97 | 73 | 16.82 | 22 | 6.34 | 38 | 8.9 |
| Resilience | Capital | 59 | 9.1 | 41 | 5.77 | 80 | 17.32 | 138 | 12.68 | 21 | 5.34 | 125 | 16.76 | 6 | 1.38 | 26 | 7.49 | 85 | 19.91 |
| | City | 65 | 10.03 | 28 | 3.94 | 65 | 14.07 | 208 | 19.12 | 40 | 10.18 | 105 | 14.08 | 77 | 17.74 | 19 | 5.48 | 21 | 4.92 |
| | Town | 121 | 18.67 | 121 | 17.04 | 122 | 26.41 | 296 | 27.21 | 55 | 13.99 | 277 | 37.13 | 61 | 14.06 | 53 | 15.27 | 60 | 14.05 |
| | Countryside | 392 | 60.49 | 279 | 39.3 | 187 | 40.48 | 433 | 39.8 | 207 | 52.67 | 230 | 30.83 | 286 | 65.9 | 204 | 58.79 | 251 | 58.78 |
| | Farmland | 11 | 1.7 | 241 | 33.94 | 8 | 1.73 | 13 | 1.19 | 70 | 17.81 | 9 | 1.21 | 4 | 0.92 | 45 | 12.97 | 10 | 2.34 |
| Having children | Yes | 377 | 58.18 | 377 | 53.1 | 236 | 51.08 | 527 | 48.44 | 226 | 57.51 | 392 | 52.55 | 242 | 55.76 | 184 | 53.03 | 256 | 59.95 |
| | No | 271 | 41.82 | 333 | 46.9 | 226 | 48.92 | 561 | 51.56 | 167 | 42.49 | 354 | 47.45 | 192 | 44.24 | 163 | 46.97 | 171 | 40.05 |

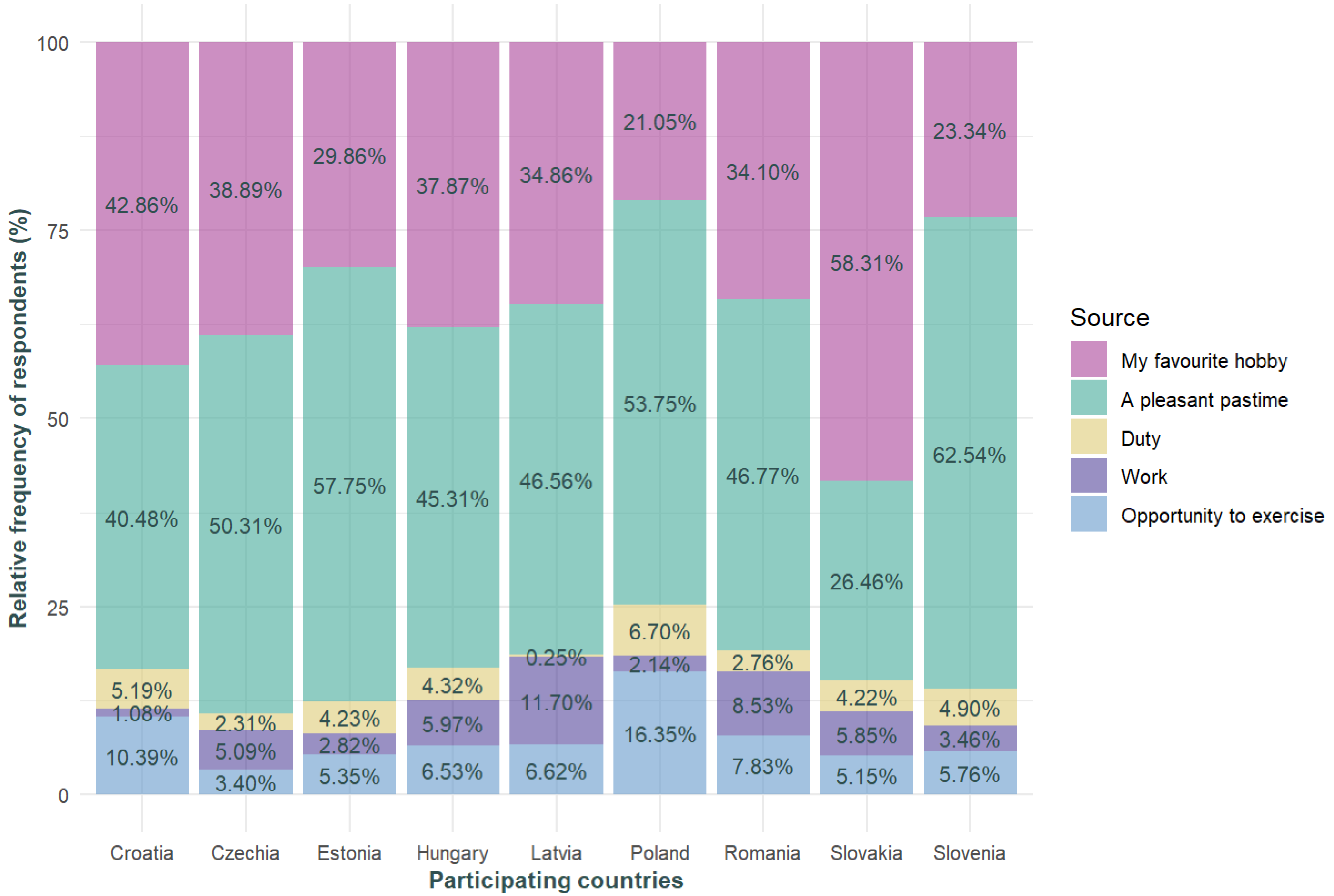


**Supplementary Material Fig. 2**: The relative frequency (%) of how respondents (n = 5255) perceived gardening in the nine participating countries.

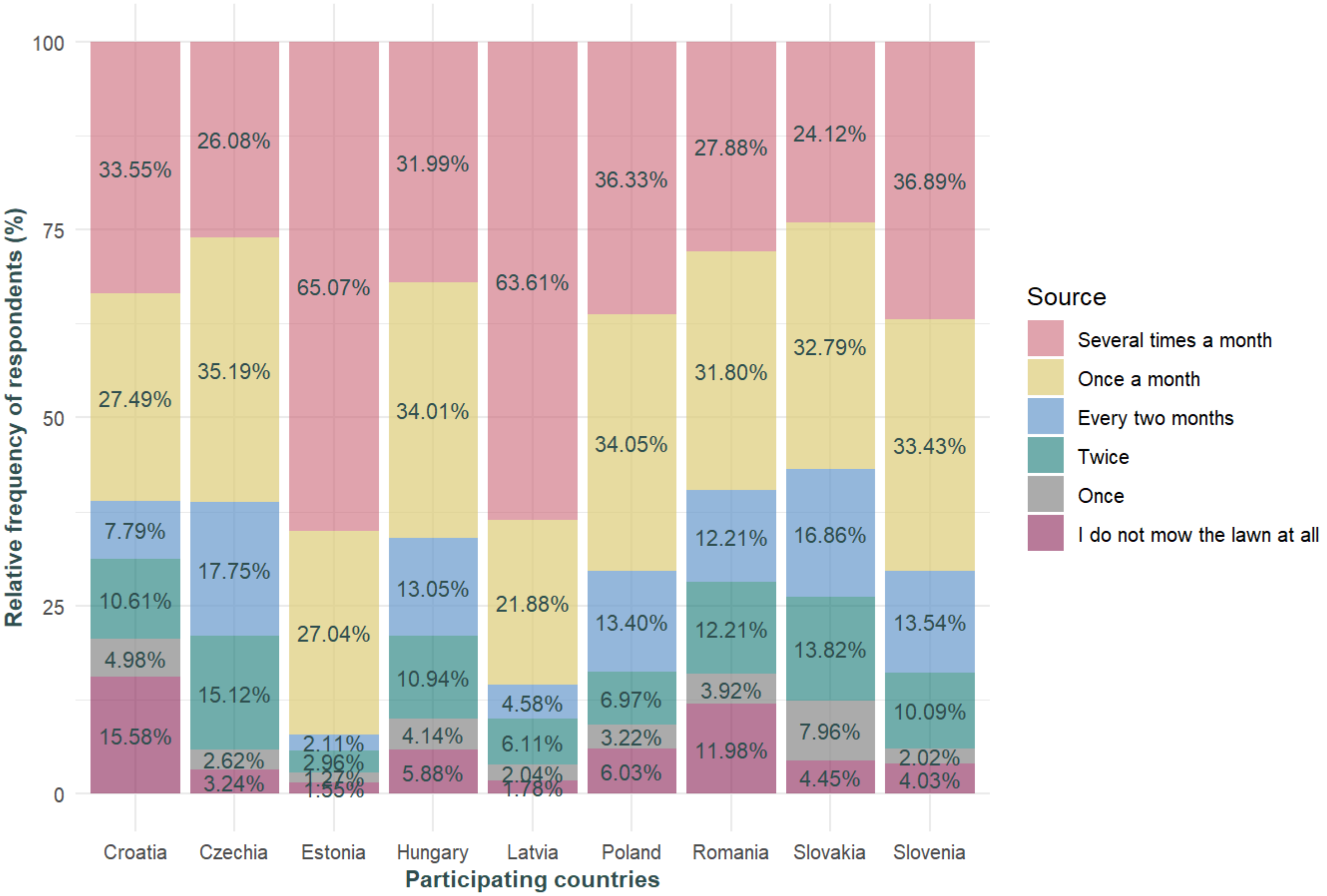


**Supplementary Material Fig. 3:** The relative frequency (%) of how often respondents (n = 5255) mowed their lawns during the growing season (between April and September) in the nine participating countries.

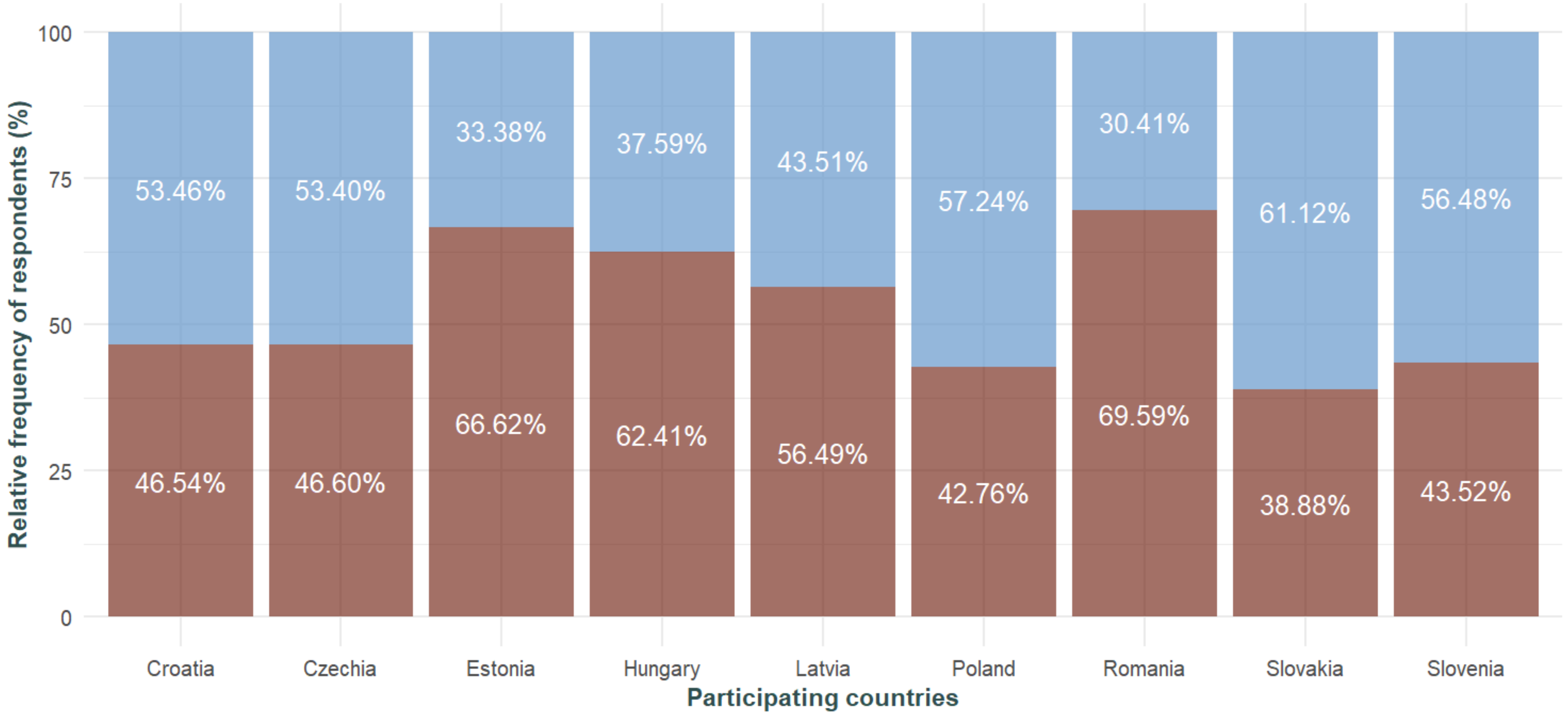


**Supplementary Material Fig. 4:** Relative frequency (%) of domestic garden owners using or not using pesticides. The response is colour-coded as follows: brown – Yes, blue – No. Pesticide types are not distinguished in these figures. If garden owners choose 'Yes', that can equally mean that they use only synthetic products, only eco or home-made pesticides, or a combination of these.

### Indices

**Supplementary Material Table 3:** Quantile values for the garden (GAR), respondents (RES), and pesticide (PES) index.

| Country | GAR | RES | PES |
|---|---|---|---|
| 0% | 14.97 | 9.25 | 24.87 |
| 25% | 47.58 | 52.17 | 60.87 |
| 50% | 56.49 | 63.25 | 76.33 |
| 75% | 64.54 | 72.67 | 97.00 |
| 100% | 91.32 | 96.75 | 100.00 |

**Supplementary Material Table 4:** The mean (range) of the tree indices (GAR – garden, RES – respondents, and PES – pesticide index) in the nine participating countries.

| Country | GAR | RES | PES |
|---|---|---|---|
| Czechia | 85.68 (25.01, 91.32) | 68.25 (26.42, 92.00) | 79.30 (33.20, 100) |
| Estonia | 58.01 (19.69, 89.94) | 61.73 (16.92, 94.50) | 74.86 (27.74, 100) |
| Croatia | 53.84 (23.08, 85.13) | 54.54 (16.83, 87.00) | 79.80 (35.12, 100) |
| Hungary | 54.96 (14.97, 90.57) | 65.69 (15.92, 95.67) | 72.20 (24.87, 100) |
| Latvia | 59.19 (25.35, 85.82) | 62.73 (23.50, 96.00) | 77.35 (39.02, 100) |
| Poland | 51.79 (15.47, 88.03) | 51.90 (9.25, 84.83) | 82.57 (36.70, 100) |
| Romania | 50.98 (21.06, 84.56) | 60.55 (14.75, 95.00) | 71.18 (31.57, 100) |
| Slovenia | 57.90 (33.84, 81.87) | 65.33 (30.75, 92.00) | 83.44 (31.80, 100) |
| Slovakia | 60.03 (23.95, 91.14) | 64.85 (16.50, 96.75) | 83.31 (40.61, 100) |

**Supplementary Material Table 5:** Pairwise comparisons of (a) GAR, (b) RES, and (c) PES indices among the nine participating countries using Wilcoxon rank sum tests. Asterisks indicate significant relationships.

| (a) | CZ | EE | HR | HU | LV | PL | RO | SI |
|---|---|---|---|---|---|---|---|---|
| EE | 0.46775 | - | | | | | | |
| HR | 4.9e-10* | 1.9e-08* | - | | | | | |
| HU | 4.9e-08* | 2.4e-06* | 0.06814 | - | | | | |
| LV | 0.58091 | 0.24071 | 4.9e-10* | 1.2e-07* | - | | | |
| PL | < 2e-16* | < 2e-16* | 0.00839* | 1.2e-07* | < 2e-16* | - | | |
| RO | < 2e-16* | < 2e-16* | 0.00051* | 1.4e-08* | < 2e-16* | 0.25489 | - | |
| SI | 0.25225 | 0.59532 | 3.2e-06* | 0.00079* | 0.11878 | 1.5e-13* | 2.1e-14* | - |
| SK | 0.05325 | 0.01050* | 2.2e-14* | 3.0e-12* | 0.19602 | < 2e-16* | < 2e-16* | 0.00296* |

| (b) | CZ | EE | HR | HU | LV | PL | RO | SI |
|---|---|---|---|---|---|---|---|---|
| EE | < 2e-16* | - | | | | | | |
| HR | < 2e-16* | < 2e-16* | - | | | | | |
| HU | 0.00350* | 1.5e-10* | < 2e-16* | - | | | | |
| LV | 1.6e-09* | 0.26033 | 7.4e-15* | 0.00014* | - | | | |
| PL | < 2e-16* | < 2e-16* | 0.00776* | < 2e-16* | < 2e-16* | - | | |
| RO | < 2e-16* | 0.31331 | 1.2e-09* | 4.7e-10* | 0.06656 | < 2e-16* | - | |
| SI | 4.7e-05* | 0.00041* | < 2e-16* | 0.09230 | 0.03953* | < 2e-16* | 8.4e-05* | - |
| SK | 1.9e-05* | 0.00015* | < 2e-16* | 0.07002 | 0.03980* | < 2e-16* | 4.2e-05* | 0.87783 |

| (c) | CZ | EE | HR | HU | LV | PL | RO | SI |
|---|---|---|---|---|---|---|---|---|
| **EE** | 0.00014* | - | | | | | | |
| **HR** | 0.85028 | 0.00029* | - | | | | | |
| **HU** | 1.3e-08* | 0.00270* | 4.3e-08* | - | | | | |
| **LV** | 0.00556* | 0.20356 | 0.01253* | 0.00162* | - | | | |
| **PL** | 0.00485* | 3.9e-14* | 0.00324* | < 2e-16* | 1.4e-08* | - | | |
| **RO** | 9.4e-09* | 0.00162* | 1.4e-08* | 0.64147 | 0.00015* | < 2e-16* | - | |
| **SI** | 0.01438* | 6.3e-12* | 0.00475* | 2.6e-15* | 6.9e-08* | 0.97237 | < 2e-16* | - |
| **SK** | 0.00162* | 5.5e-13* | 0.00075* | < 2e-16* | 3.1e-09* | 0.37674* | < 2e-16* | 0.36218 |

### GBM modelling

**Supplementary Material Table 5**: Relative influence of factors generated from the Gradient Boosting Machine (GBM) model (R-squared: 0.12, RMSE: 11.29) for predicting garden (GAR) index.

| Variables | Relative influence |
|---|---|
| Country | 26.715 |
| Average time spent with gardening | 17.280 |
| Age | 11.625 |
| Gardening perception | 11.400 |
| Longitudes | 9.167 |
| Gardening experience | 7.640 |
| Latitudes | 6.030 |
| PPS | 4.667 |
| Education level | 4.454 |
| Having children | 0.823 |
| Gender | 0.198 |

**Supplementary Material Table 6**: Relative influence of factors generated from the Gradient Boosting Machine (GBM) model (R-squared: 0.24, RMSE: 13.13) for predicting respondents (RES) index.

| Variables | Relative influence |
|---|---|
| Country | 39.54 |
| Gardening perception | 24.51 |
| Average time spent with gardening | 12.16 |
| Age | 7.93 |
| Gardening experience | 4.52 |
| Longitudes | 3.97 |
| Latitudes | 3.68 |
| PPS | 1.50 |
| Education level | 1.09 |
| Gender | 0.61 |
| Having children | 0.49 |

**Supplementary Material Table 7**: Relative influence of factors generated from the Gradient Boosting Machine (GBM) model (R-squared: 0.08, RMSE: 18.19) for predicting pesticide (PES) index.

| **Variables** | **Relative influence** |
|---|---:|
| Country | 46.10 |
| Average time spent with gardening | 16.94 |
| Gardening experience | 12.06 |
| Gardening perception | 6.69 |
| Longitudes | 4.59 |
| Latitudes | 4.15 |
| Education level | 3.15 |
| Age | 2.83 |
| Gender | 2.66 |
| Having children | 0.84 |
| PPS | 0.00 |

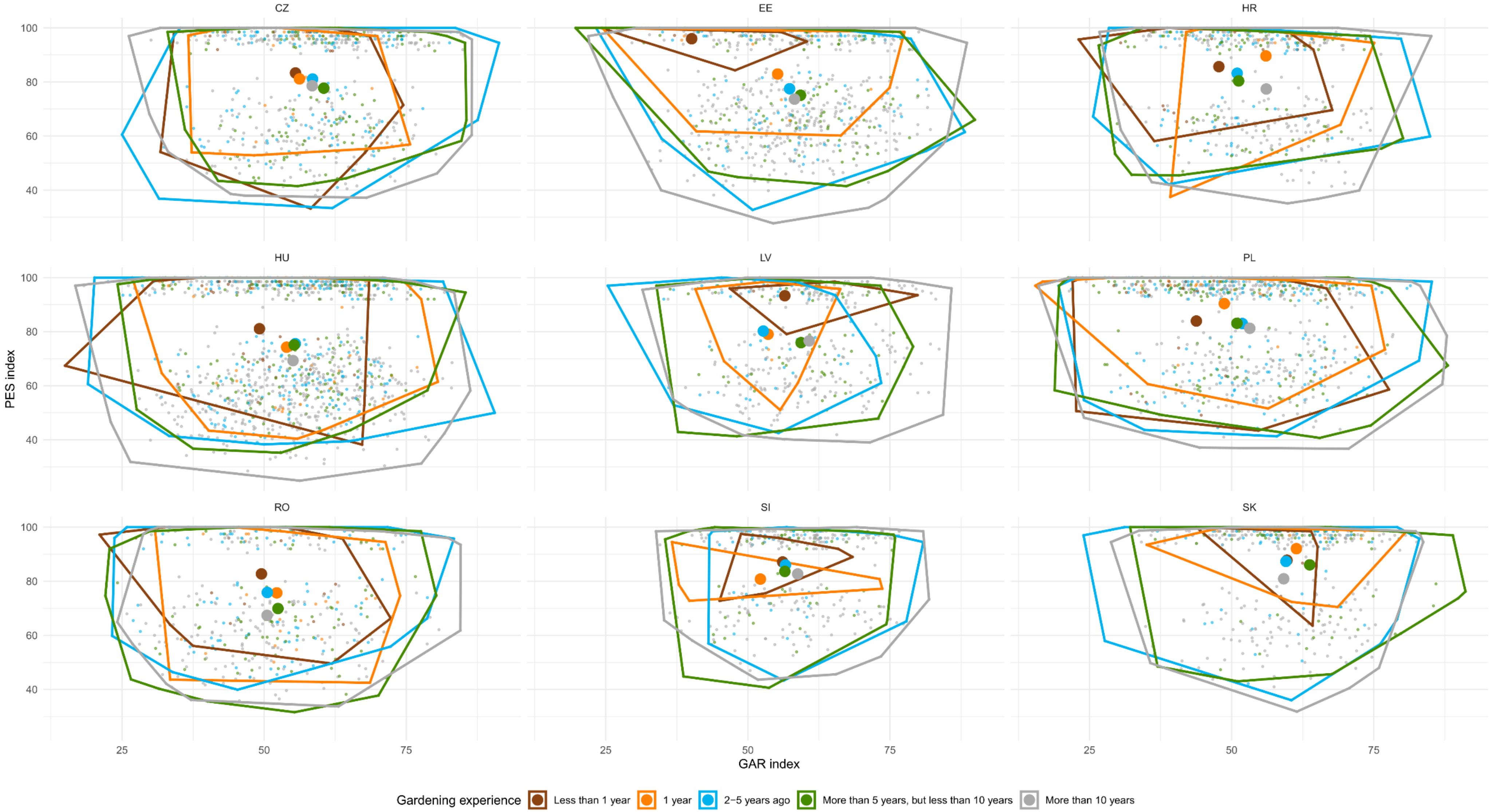


**Supplementary Material Fig. 5**: The GAR index as a function of the PES index. Each transparent point represents a respondent, coloured according to how long he/she has been gardening. The response is colour-coded as follows: brown – less than one year, orange – one year, light blue – two to five years, green – more than five, but less than ten years, grey – more than ten years. Each large point represents the mean of the responses.

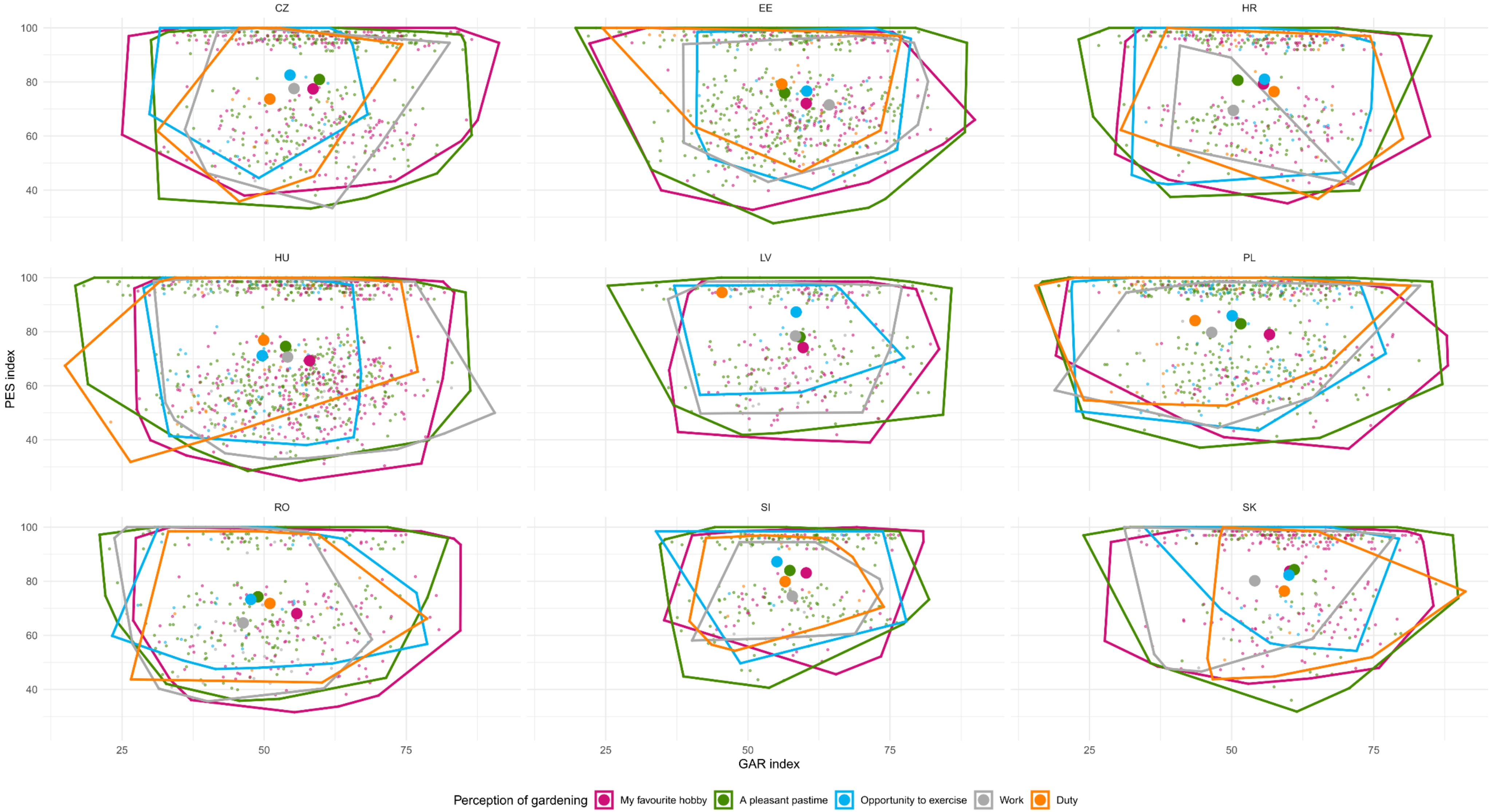


**Supplementary Material Fig. 6**: The GAR index as a function of the PES index. Each transparent point represents a respondent, coloured according to how garden owners perceive gardening. The response is colour-coded as follows: pink – a favourite hobby, green – a pleasant pastime, blue – an opportunity to exercise, grey – duty, orange – work. Each large point represents the mean of the responses.

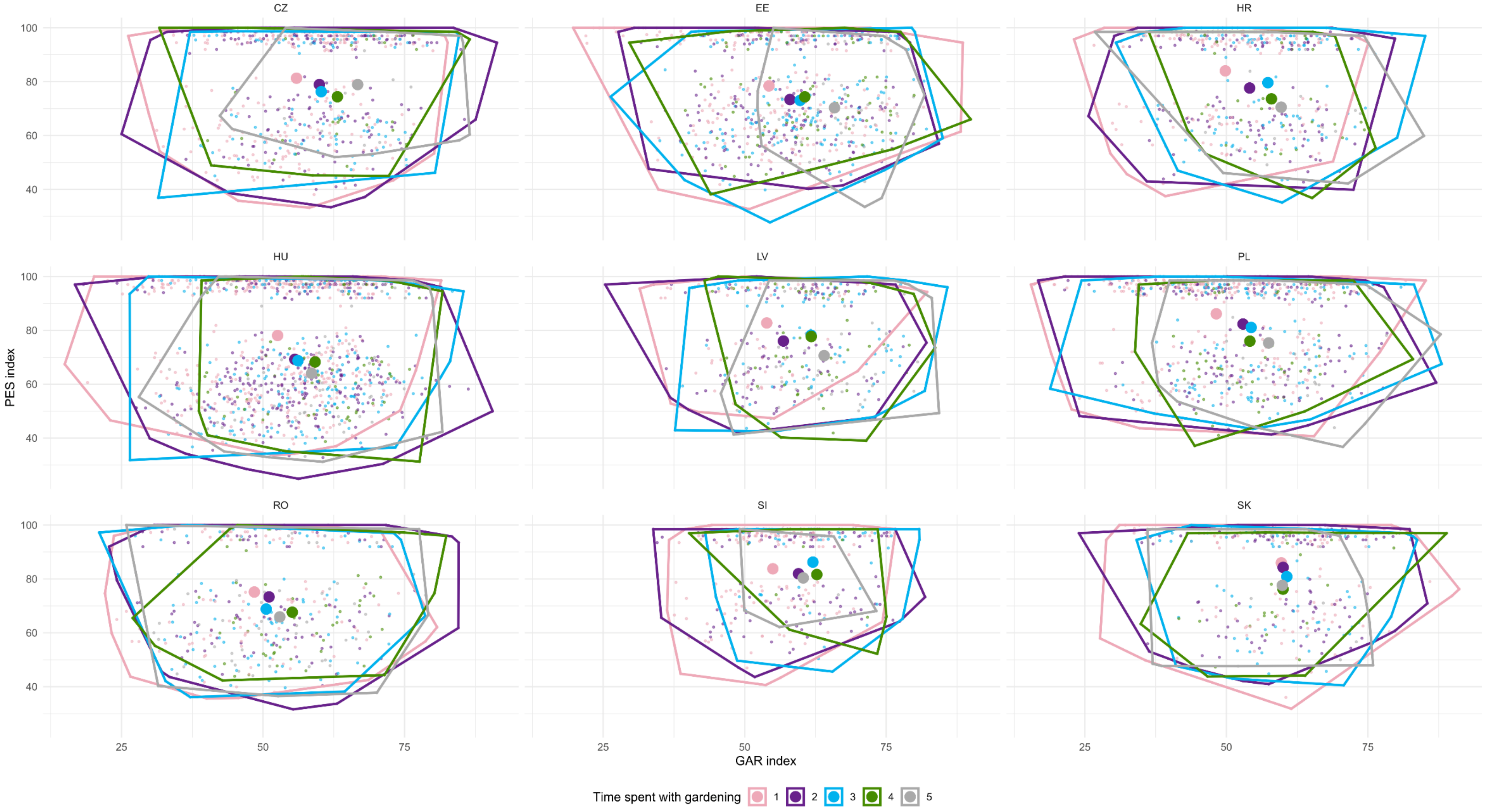


**Supplementary Material Fig. 7:** The GAR index as a function of the PES index. Each transparent point represents a respondent, coloured according to during the growing season (April to September), on an average day, approximately how many hours respondents spend with gardening. Time spent with gardening is shown in a self-reported scoring system as indicated by the respondents. The response is colour-coded from one to two hours per day (light-pink) as the minimum, to the even twelve hours/day (light-grey) as the maximum hours spent with gardening.

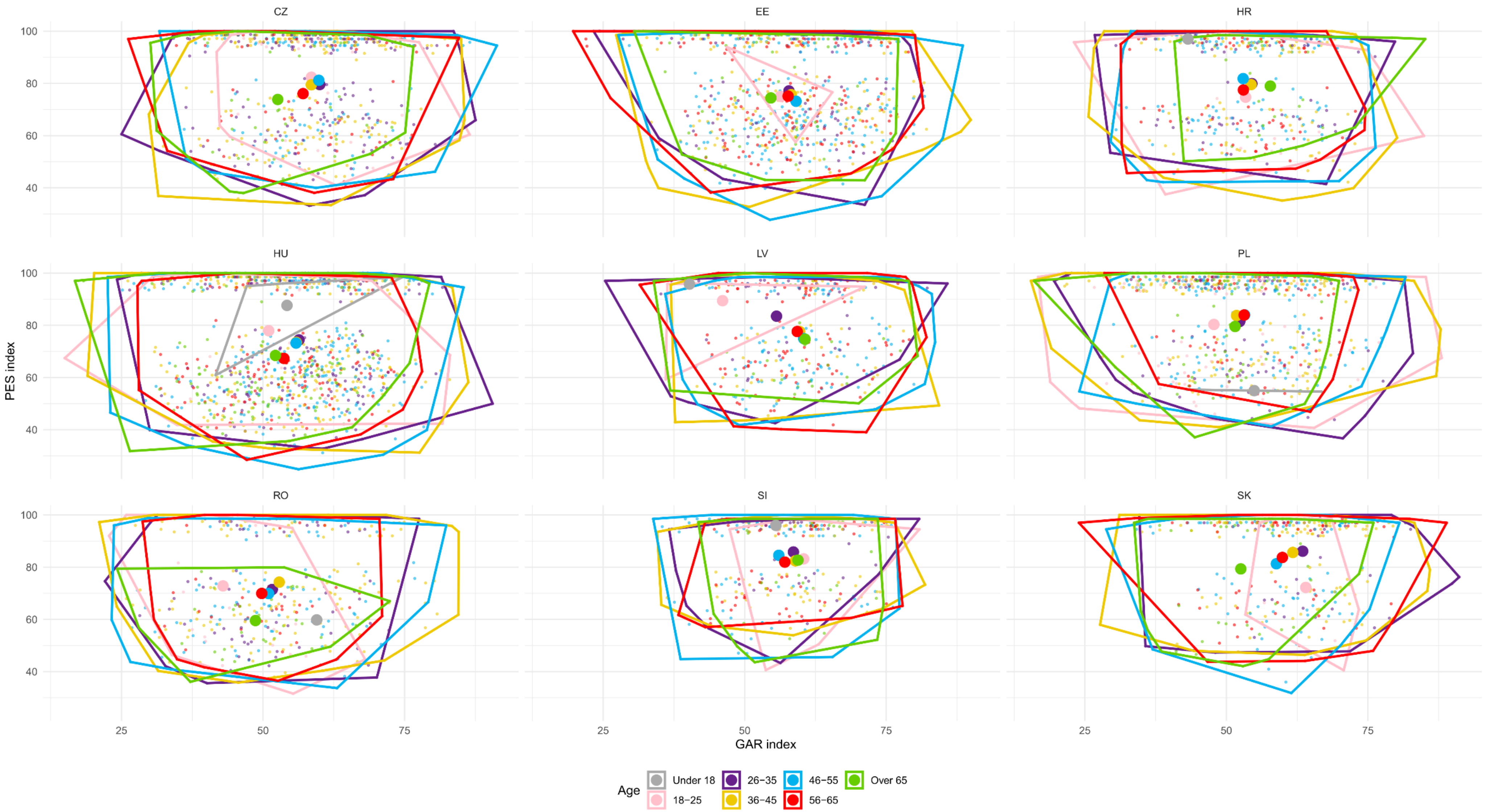


**Supplementary Material Fig. 8:** The GAR index as a function of the PES index. Each transparent point represents a respondent, coloured according to the age groups. The response is colour-coded as follows: grey – age under 18, pink – age 18-25, purple – age 26-35, yellow – age 36-45, blue – age 46-56, red – age 56-65, green – age over 65. Each large point represents the mean of the responses.

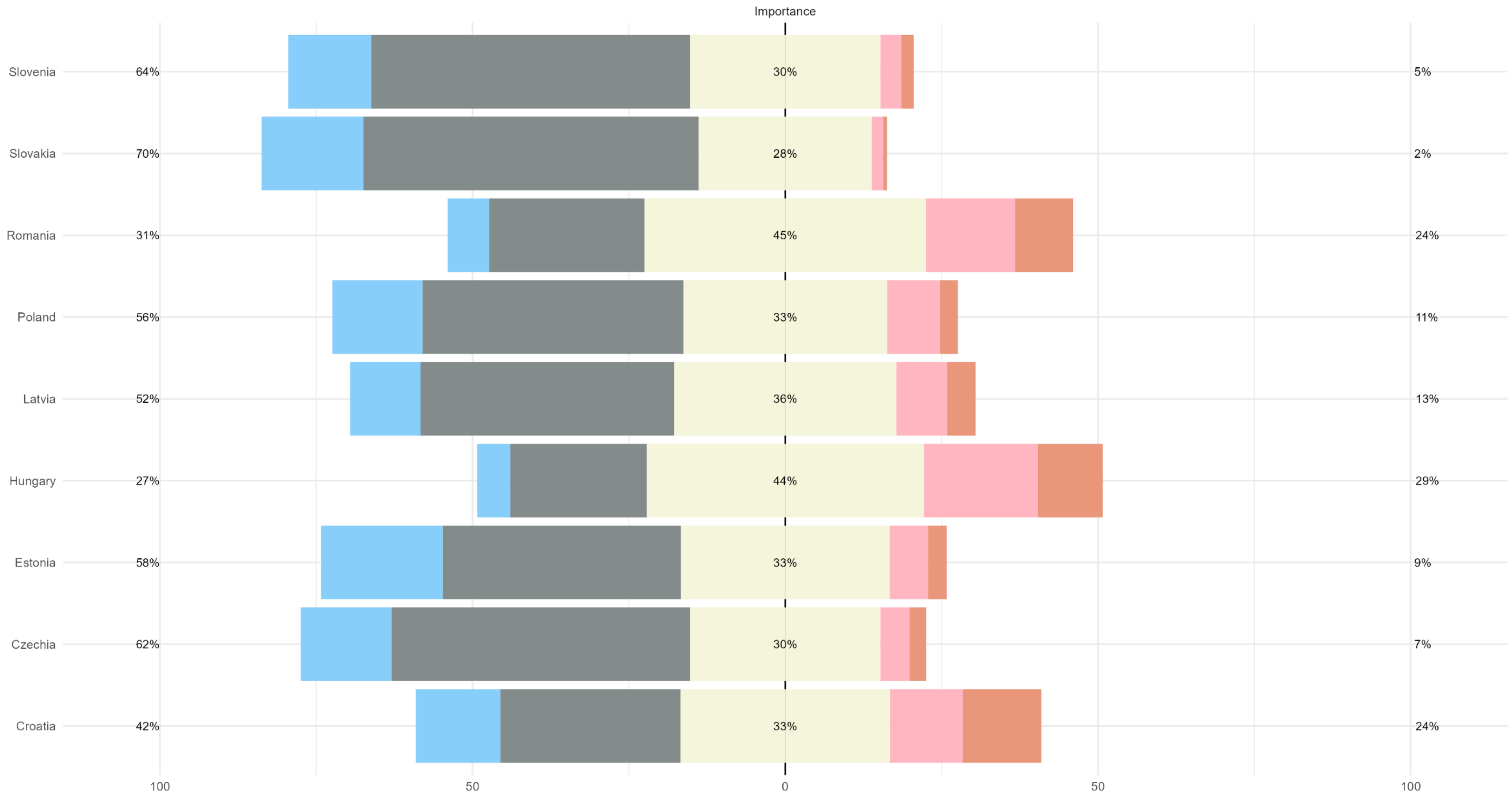


**Supplementary Material Fig. 9:** Relative frequency (%) of the considered importance of pesticide use in gardens according to those garden owners who use pesticides (n = 2 829). The response is colour-coded as follows: light blue –not at all, grey – negligible, light yellow – moderately important, light pink – important, dark orange – crucial.

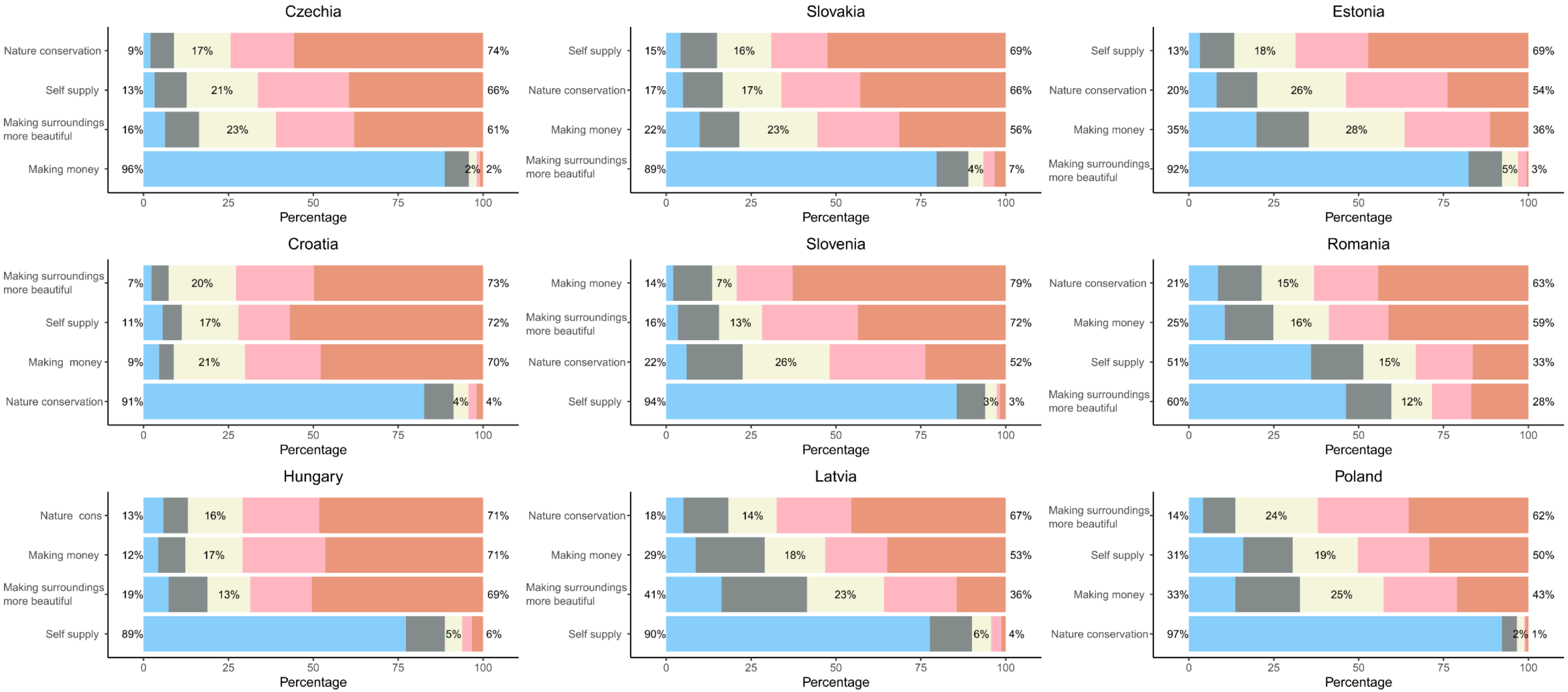


**Supplementary Material Fig. 10**: Relative frequency (%) of respondents' opinions on what extent the listed aspects influence their gardening habits. The response is colour-coded as follows: light blue – it does not affect at all, grey – negligible, light yellow – moderately, light pink – important, dark orange – this is the most important. For each country, the most important aspect for the most respondents is ranked first, and the least important aspect for the most respondents is ranked last.
