## Supplementary Methods for "Improving biodiversity in Central and Eastern European domestic gardens needs regionally scaled strategies"

**This file includes:**

Explanation of the scoring system

Supplementary Methods Table 1-3

Explanation of indices components

Additional information

References

### 1. Explanation of the scoring system

We classified the gardens, the garden owners' attitude and motivation for gardening, as well as, their pesticide use through an answer-based scoring system. As a result, each respondent got three points, one for each index. The scoring scale was between 0 and 100 points for each index.

- The garden (GAR) index indicates how high diversity the garden can potentially support.
- The respondent (RES) index reflects on garden owners' attitude to maintaining/creating a biodiverse garden.
- The pesticide (PES) index indicates the degree of pesticide load in gardens.

#### 1.1 Garden (GAR) index

##### 1.1.1 Components of the GAR index

**Table 1**: The components of the scoring system for the garden (GAR) index. The number in the last column shows the weight (%) with which the respective index component contributed into the index.

| Index component | Question(s) | Answer options | Answers' scores | Answers' maximum values | Formula for calculating | Weight (%) |
|---|---|---|---|---|---|---|
| 1 | What is the area of your garden? | Under 10 $m^2$ | 1 | 5 | Equals to score (no calculation) | 10 |
| | | 10-50 $m^2$ | 2 | | | |
| | | 50-100 $m^2$ | 3 | | | |
| | | 100-500 $m^2$ | 4 | | | |
| | | Over 500 $m^2$ | 5 | | | |
| 2 | What type of land your garden is adjacent to? † | Urban area | 0 | 16 | Sum of scores | 20 |
| | | Garden, park | 2 | | | |
| | | Agricultural area/farm | 2 | | | |
| | | Forest | 4 | | | |
| | | Field, meadow | 3 | | | |
| | | Wetland | 4 | | | |
| | | Other | 1 | | | |
| 3 | How large area do the listed (*Vegetables; Herbs; Fruit trees; Grapes; Ornamental plants/trees; Evergreens; Lawn; Undisturbed area/fallow*) plants | Do not have | 0 | 10.20444 | $GSw(\theta)=\sum_i w_i p_i (1-p_i)$ © | 20 |
| | | Very small | 1 | | | |
| | | Medium | 2 | | | |

| | | | | | | |
|---|---|---|---|---|---|---|
| | take up in your garden? *(Weight of plants type: 1.5, 1.5, 1.75, 0.5, 1, 0.5, 0.5, 2)* | Significant<br>Most of it | 3<br>4 | | GSw(θ)length(*chosen answer*[*if the value not 0*]) | |
| 4 | Do you have a pond in your garden? | Yes<br>No | 0<br>3 | 6 | Sum of scores | 10 |
| | Do you leave unmown patches when you mow? | Yes<br>No | 0<br>1 | | | |
| | Do you have any undisturbed areas in your garden? | Yes<br>No | 0<br>2 | | | |
| 5 | How often do you mow the lawn during the growing season? | Several times a month<br>Once a month<br>Every two months<br>Twice<br>Once<br>I do not mow the lawn at all | 0<br>1<br>2<br>3<br>4<br>5 | 5 | Equals to score (no calculation) | 5 |
| 6 | Do you use artificial grass in your garden? | Yes*<br>No | -1<br>0 | 0 | Equals to score (no calculation) | 5 |
| 7 | Do you use herbicides in your garden? | Yes*<br>No | -1<br>0 | 0 | Equals to score (no calculation) | 5 |
| 8 | Do you support beneficial insects to promote the biocontrol of weeds and pests? | Yes<br>No | 1<br>0 | 1 | Equals to score (no calculation) | 5 |

| | | | | | | |
|---|---|---|---|---|---|---|
| 9 | What pollinators do you regularly observe/see in your garden? † | Butterflies | 1 | 9 | Number of ticks | 10 |
| | | Bumblebees | 1 | | | |
| | | Honeybees | 1 | | | |
| | | Other wild bees | 1 | | | |
| | | Wasps | 1 | | | |
| | | Ants | 1 | | | |
| | | Hoverflies | 1 | | | |
| | | Other flies | 1 | | | |
| | | Beetles | 1 | | | |
| | | None of the above | 0 | | | |
| 10 | How do you support wild pollinators? † | Creating/preserving natural habitats | 3 | 3 | Number of ticks | 10 |
| | | Creating artificial habitats | 2 | | | |
| | | Providing food sources | 2 | | | |
| | | Water sources | 1 | | | |
| | | I do not support them actively | 0 | | | |

* The original categorical replies were re-categorised for analytical purposes.

† Multiple choices were allowed.

© Citation for the Weighted Gini-Simpson Index: https://www.hindawi.com/journals/ijecol/2012/478728/

##### 1.1.2 Explanation of the components of the GAR index

1. The larger-sized gardens scored higher because they are more likely to maintain a higher biodiversity (Burks & Philpott, 2017; Daniels & Kirkpatrick, 2006; Delahay et al., 2023; Fontaine et al., 2016; Quistberg et al., 2016).
2. Each domestic garden received a summarised score based on its adjacency to different habitats. Urban areas score zero points, indicating them as being the least favourable. Wetlands and forests scored the highest, as they can support diverse communities of living organisms, benefiting the domestic gardens' biodiversity (Braschler et al., 2020).

3. We evaluated the size and variety of plant types in domestic gardens. Vines, evergreens, and lawns received lower scores due to factors like pesticide use, limited insect attraction (especially to evergreens), and the predominant grass cover (usually without any flowering plants). Fallow/undisturbed areas received the highest score. The Weighted Gini-Simpson Index© was calculated based on the number and relative importance of different habitat types in the garden.
4. Ponds received the highest score (3 points) for promoting/supporting biodiversity. Undisturbed areas received a higher score (2 points) than the unmown areas (1 point) because an unmown area does not necessarily imply that it is always undisturbed, but undisturbed areas are always left unmown. Therefore, being unmown implies less reduction in disturbance. The scores from these components are summed, with a maximum reachable score of 6.
5. Reduced mowing frequency results in a higher score, as it helps to create and preserve/support high diversity in gardens (Sehrt et al., 2020; Wintergerst et al., 2021).
6. The use of artificial grass was considered negative because it is unsuitable as a habitat and it has negative impacts on the environment, including microplastic pollution (Francis, 2018; Sánchez-Sotomayor et al., 2023).
7. We considered herbicide use as strongly harmful as it extirpates wildflowers essential for attracting pollinators and herbicides also have direct negative effects on pollinators (Tassin de Montaigu & Goulson, 2023).
8. Supporting ecosystem service provider insects, like ladybirds, earwigs, and hoverflies contributes to biodiversity and encourages conservation actions (Gardiner et al., 2014).
9. Points were awarded for each pollinator group observed in the garden, as a direct indicator of species richness.
10. Creating and preserving natural habitats received the highest score, providing the most significant support for biodiversity.

#### 1.2 Respondents (RES) index

##### 1.2.1 Components of the RES index

**Table 2:** The components of the scoring system for the respondent (RES) index. The number in the last column shows the weight (%) with which each index component contributed into the index.

| Index component | Question(s) | Answer options | Answers' scores | Answers' maximum values | Formula for calculating value | Weight (%) |
|---|---|---|---|---|---|---|
| 1 | To what extent do the '*Nature conservation*' influence your gardening habit? | 1 (It does not affect at all)<br>2<br>3<br>4<br>5 (This is the most important) | 1,<br>2,<br>3,<br>4,<br>5 | 5 | Equals to score (no calculation) | 15 |
| 2 | Plants/Birds/Insects of my garden… (please, finish the sentences) | I know.<br>I know, and I explore them consciously.<br>I do not know them well but I am trying to get to know it better.<br>I do not know, but I want to know.<br>I do not know. | 3,<br>4,<br>2,<br>1,<br>0 | 12 | Number of ticks | 13 |
| 3 | Do you support wild pollinators? | Yes*<br>No | 0,<br>1 | 2 | No = 0, 1 support type = 1, multiple support type = 2 | 12 |

| | | | | | | |
|---|---|---|---|---|---|---|
| 4 | Do you think your garden is pollinator-friendly? | Yes<br>No | 1,<br>0 | 1 | Equals to score (no calculation) | 5 |
| 5 | Can you imagine your garden being part of a garden network that helps maintain biodiversity? | Yes<br>Maybe<br>No | 2,<br>1,<br>0 | 2 | Equals to score (no calculation) | 15 |
| 6 | Do you usually submit data on arthropods in your garden to citizen science-based biodiversity recording platforms? | 1 (No)<br>2<br>3<br>4<br>5 (Yes, several times a week) | 0,<br>1,<br>2,<br>3,<br>4 | 4 | Equals to score (no calculation) | 10 |
| 7 | Have you heard about the #NoMowMay 2022 campaign and if so, have you joined it? | Yes, and I did not mow in May.<br>Yes, but I did mow in May.<br>I have not heard about this campaign. | 1,<br>-1,<br>0 | 1 | Equals to score (no calculation) | 5 |
| 8 | To what level do you think synthetic pesticides can threaten invertebrates? | Highly<br>Significantly<br>Moderately<br>Negligibly<br>Not et all | 3,<br>3,<br>2,<br>1,<br>0 | 3 | Equals to score (no calculation) | 3 |
| 9 | How do you learn/gather knowledge/information about gardening? † | Gardening journals/magazines, books<br>Internet (e.g. gardening forums)<br>Social media platforms (e.g. Facebook groups)<br>Self-training groups<br>Consulting with professional gardeners/agronomists | 1,<br>1,<br>1,<br>1,<br>1, | 3 | if length(answers) = 1 → 1 point;<br>if length(answers) = 2, 3 or 4 → 2 points;<br>if length(answers) >= 5 → 3 points | 12 |

| | | | | | | |
|---|---|---|---|---|---|---|
| | | TV, radio | 1, | | | |
| | | Other | 1 | | | |
| 10 | Do you document the development/changes of your garden with pictures, videos, and/or notes? | Yes<br>No | 1,<br>0 | 1 | Equals to score (no calculation) | 10 |

* The original categorical replies were re-categorised for analytical purposes.

† Multiple choices were allowed.

##### 1.2.2 Explanation of the components of the RES index

1. Garden owners who strongly prioritise nature conservation in their gardening habits contribute positively to their garden's biodiversity. Therefore, the higher they value conservation, the higher scores they receive.
2. We awarded the highest scores to responses indicating an active and conscious effort to learn about the plants, birds, and insects in domestic gardens.
3. The question 'Do you support wild pollinators?' was also included in the diversity (DIV) index, but in this (RES) index, we based scoring solely on respondents' intentions to support wild pollinators.
4. While this question is theoretical, it is relevant to the index and indicates the garden owner's understanding of what it means to be 'pollinator-friendly'. The ambition to create a pollinator-friendly domestic garden is particularly important for supporting biodiversity.
5. Despite being theoretical, this question holds significant importance in our scoring system, particularly for mapping the availability of domestic gardens for a high-diversity garden network. The ambition of garden owners to maintain high diversity is necessary.
6. Domestic garden owners who utilise citizen science-based biodiversity recording platforms, such as iNaturalist, are likely to be more open to preserving biodiversity.
7. The 'No Mow May' campaign (*Plantlife's No Mow May Movement*, 2024) did not reach all participating countries equally, but if someone was aware of the campaign but did not participate, we assigned a lower score.
8. Garden owners' perceptions of the harmful effects of synthetic pesticides on invertebrates may influence how consciously they use these chemicals in their gardens.

9. Gathering gardening information from multiple channels enhances confidence in gardeners' access to a wide range of perspectives. Those relying on a single information source receive fewer points, as it carries the risk of incomplete information, such as the importance of biodiversity conservation.
10. Documenting the garden's development and changes is a positive activity as it promotes conscious gardening practices.

#### 1.3 Pesticide (PES) index

##### 1.3.1 Components of the RES index

**Table 3:** The components of the scoring system for the pesticide (PES) index. The number in the last column shows the weight (%) with which each index component contributed into the index.

| Index component | Question(s) | Answer options | Answers' scores | Answers' maximum values | Formula for calculating value | Weight (%) |
|---|---|---|---|---|---|---|
| 1 | Do you regularly fertilizer your garden? † | I use synthetic fertilizer | 2 | 4 | Number of ticks | 6 |
| | | I use animal manure | 1 | | | |
| | | I use compost | 1 | | | |
| | | I do not fertilize | 0 | | | |
| 2 | Do you have a dog(g) and/or cat(s) in your garden? If so, do you regularly treat it/them against fleas in the form of preventive collars or drops? | I have it/them, and I treat it/them several times a year | 2 | 4 | Number of ticks | 5 |
| | | I have it/them, and I treat it/them once a year | 1 | | | |
| | | I have it/them, but I do not give it/them preventive treatments | 0 | | | |
| | | I do not have it/them | 0 | | | |
| 3⁑ | What type of pesticides(s) do you use? † | Conventional pesticides/synthetic pesticides | 4 | 7 | Number of ticks | 18 |
| | | Eco/Bio/Green labelled pesticides | 2 | | | |
| | | Self-made pesticides, home practices | 1 | | | |

| | | | | | | |
|---|---|---|---|---|---|---|
| 4[*] | How many different pesticides do you use in your garden? | Only 1 | 1 | 4 | Equals to score (no calculation) | 14 |
| | | 2-4 | 2 | | | |
| | | 5-10 | 3 | | | |
| | | More than 10 | 4 | | | |
| 5[*] | Do you use additives with pesticides? | Yes | 1 | 1 | Equals to score (no calculation) | 5 |
| | | No | 0 | | | |
| 6[*] | What kind of organisms do you use pesticides against in your garden?[†] | Arthropods | 3 | 20 | Number of ticks | 14 |
| | | Snails | 2 | | | |
| | | Nematodes | 2 | | | |
| | | Vertebrates | 2 | | | |
| | | Fungi | 1 | | | |
| | | Bacteria | 1 | | | |
| | | Viruses | 1 | | | |
| | | Weeds | 3 | | | |
| | | Mooses | 2 | | | |
| | | Trees | 2 | | | |
| | | Other | 1 | | | |
| 7[*] | How important the *risk on bees* is for you when you choose a pesticide? | Crucial | 0 | 4 | Equals to score (no calculation) | 6 |
| | | Important | 1 | | | |
| | | Moderately | 2 | | | |
| | | Negligible importance | 3 | | | |
| | | Not important | 4 | | | |
| 8[*] | How important the *risk on human* is for you when you choose a pesticide? | Crucial | 0 | 4 | Equals to score (no calculation) | 6 |
| | | Important | 1 | | | |
| | | Moderately | 2 | | | |

| | | | | | | |
|---|---|---|---|---|---|---|
| | | Negligible importance | 3 | | | |
| | | Not important | 4 | | | |
| 9‡ | Where do you usually buy the pesticides? † | Horticulture/floral shops | 1 | 15 | Number of ticks | 6 |
| | | Agricultural shops | 2 | | | |
| | | Supermarkets or other big shops | 3 | | | |
| | | Online shops | 3 | | | |
| | | From acquaintances | 4 | | | |
| | | Other | 2 | | | |
| 10‡ | Do you read the direction for use on the packaging? | Yes, before I buy | 1 | 5 | Equals to score (no calculation) | 6 |
| | | Yes, the first time I use | 3 | | | |
| | | Yes, whenever I use | 2 | | | |
| | | Occasionally, if I do not forget | 4 | | | |
| | | Rarely or never | 5 (or 0[i]) | | | |
| 11‡ | How important is pesticide use in your garden? | 1 (Not at all) | 1 | 5 | Equals to score (no calculation) | 14 |
| | | 2 | 2 | | | |
| | | 3 | 3 | | | |
| | | 4 | 4 | | | |
| | | 5 (Crucial) | 5 | | | |

‡ Questions for only pesticide users.

† Multiple choices were allowed.

i In the case when someone only uses '*Self-made pesticides, home practices*' and '*Rarely or never*' reads the labels 0 point was given (instead of 5 which would indicate that someone never reads the labels).

#### 1.3.2 Explanation of the components of the PES index

1. The use of synthetic fertilizer received the lowest scores, as it is an unnatural product commonly and often unnecessarily used in domestic gardens.

2. Anti-flea products for dogs and cats, usually containing neonicotinoids as its main components, contribute to pesticide pollution in the domestic garden and the environment (Perkins & Goulson, 2023; Preston-Allen et al., 2023). Residue from these products on pets can wash into the soil and freshwater (spillover).
3. Synthetic pesticides get the worst score. However, respondents who use all three types of pesticides (synthetic, eco/bio labelled, and self-made pesticides) receive the highest (worst) score.
4. The more products someone uses, the lower the score they receive, as increased product usage hampers the establishment of a biodiverse domestic garden, regardless of whether synthetic or bio products are used.
5. Additives boost the effect and improve the physical and chemical properties of pesticides.
6. We assigned different scores to organisms based on their importance in our questionnaire. Arthropods and weeds received higher points than others. The more organisms someone uses pesticides against, the higher their score.
7. Prioritizing minimizing the impact of pesticides on bees when choosing a pesticide can indicate a garden owner's environmental consciousness.
8. Garden owners who show little interest in the risks of pesticides to humans are less likely to be concerned about their impact on other organisms, such as insects.
9. Horticulture/floral shops with expert advice receive better scores, while shops with limited access to advice or aggressive advertising strategies score lower.
10. If someone only uses ‘Self-made pesticides, home practices’ as plant protection products and ’Rarely or never’ reads the labels on pesticide bottles, they receive 0 points (instead of 5, which would indicate that they never read the labels).
11. The higher the importance of pesticide use to a respondent, the higher their score.

Only respondents who do not use pesticides in any form can get zero point. Yet, the index can go up to 11 per cent even for respondents who do not use pesticides because they still can get points after two index components (fertilizer usage and pets flea treatment). For analytical purposes and the comparability with the other two indexes, we reversed the scale of PES index.

#### 2. Additional information

The original categorical replies of our questionnaire were re-categorised for analytical purposes on a few occasions. For instance, after the question about the use of artificial grass in their gardens, respondents were given ‘Yes’ or ‘No’ instead of the original three responses (‘Yes, regularly’, ‘Yes, sometimes’, ‘No’). Similarly, a binary category (‘Yes’, ’No’) was assigned to the question regarding the use of herbicides in their domestic gardens instead of the original ones (Table 1-3).

Even though **herbicides** fall under the category of pesticides, due to their significance in gardening, we distinguished them from other products, such as insecticides and fungicides.

We define '**unmown patches**' as areas of the lawn that garden owners intentionally leave uncut, while '**undisturbed areas/fallow**' denote permanently undisturbed areas unaffected by mowing. While responses regarding garden owners' familiarity with plant, insect, and bird diversity in their domestic garden were self-assessed and therefore subjective, we consistently refer to them as 'knowledge.'

We used images illustrating representative members of pollinator groups commonly found in domestic gardens on the question evaluating the garden insect pollinator diversity.

The No Mow May campaign (NMMc) (*Plantlife’s No Mow May Movement*, 2024) urged garden owners not to mow their lawns in May, as this is the month with the most abundant food sources for pollinators in the Northern Hemisphere. Given that the campaign reached several European countries (in some countries, e.g. Hungary, the campaign’s name was translated), we asked the garden owners about their awareness and participation.
